# iSMNN: Batch Effect Correction for Single-cell RNA-seq data via Iterative Supervised Mutual Nearest Neighbor Refinement

**DOI:** 10.1101/2020.11.09.375659

**Authors:** Yuchen Yang, Gang Li, Yifang Xie, Li Wang, Yingxi Yang, Jiandong Liu, Li Qian, Yun Li

**Affiliations:** Department of Pathology and Laboratory Medicine, University of North Carolina, Chapel Hill, NC 27599, USA; McAllister Heart Institute, University of North Carolina, Chapel Hill, NC 27599, USA; Department of Statistics and Operations Research, University of North Carolina, Chapel Hill, NC 27599, USA; Frontier Science Center for Immunology and Metabolism, Wuhan University, Wuhan, Hubei 430071, China; Department of Statistics, Sun Yat-sen University, Guangzhou, Guangdong 510275, China; Department of Biostatistics, University of North Carolina, Chapel Hill, NC 27599, USA; Department of Genetics, University of North Carolina, Chapel Hill, NC 27599, USA; Department of Computer Science, University of North Carolina, Chapel Hill, NC 27599, USA

## Abstract

Batch effect correction is an essential step in the integrative analysis of multiple single cell RNA-seq (scRNA-seq) data. One state-of-the-art strategy for batch effect correction is via unsupervised or supervised detection of mutual nearest neighbors (MNNs). However, both two kinds of methods only detect MNNs across batches on the top of uncorrected data, where the large batch effect may affect the MNN search. To address this issue, we presented iSMNN, a batch effect correction approach via iterative supervised MNN refinement across data after correction. Our benchmarking on both simulation and real datasets showed the advantages of the iterative refinement of MNNs on the performance of correction. Compared to popular alternative methods, our iSMNN is able to better mix the cells of the same cell type across batches. In addition, iSMNN can also facilitate the identification of differentially expression genes (DEGs) relevant to the biological function of certain cell types. These results indicated that iSMNN will be a valuable method for integrating multiple scRNA-seq datasets that can facilitate biological and medical studies at single-cell level.

## INTRODUCTION

With the rapidly improving technologies and decreasing sequencing costs, large-scale single-cell RNA-sequencing (scRNA-seq) studies examining tens of thousands to even millions of cells are becoming increasingly common [1,2]. Integrated analyses of cells across multiple studies (or batches) enable an increase in sample size for more powerful analysis and more comprehensive understanding of various biological questions and processes [3–7]. In addition, reusing published scRNA-seq datasets not only maximizes the value of existing data, but also substantially reduces the costs of generating new data. However, existing datasets may be produced across multiple time points, via different experimental protocols, and/or by various laboratories, eliciting systematic differences between different batches (also known as “batch effects”), which present grand challenges to integrative analyses across multiple datasets [8]. When not properly corrected, batch effects may lead to spurious findings and/or failure to identify biological signals including novel cell type(s) and differentially expressed genes [8–10].

To address this problem, a number of methods have recently been developed for batch effects correction [11–20]. Of them, the correction strategy based on mutual nearest neighbor (MNN) detection has been widely used in several state-of-the-art methods, such as MNNcorrect [11] and Seurat v3 [15], with a promising performance. However, these methods tend to err as once wrongly matching the near neighbors for the cells belonging to different cell types across batches (**Figure 1B and C; Supplementary Section 1; Supplementary Figure S1**). Compared to unsupervised correction methods, our previous developed method SMNN supervised searches MNNs by incorporating the cell type label information that SMNN restricts the detection of MNNs within the same cell type across batches [18]. The benchmarking results in both simulated and real datasets demonstrated that SMNN outperforms alternative unsupervised methods, MNNcorrect and Seurat v3, in terms of both reduced differences across batches and improved maintenance of cell-type specific features [18]. However, SMNN searches MNNs on the top of original expression matrices. The number of MNNs is supposed to be small when the batch effect across samples is large, which may lead to less accurate correction results compared to those with a larger amount of MNNs.

**Fig. 1.**
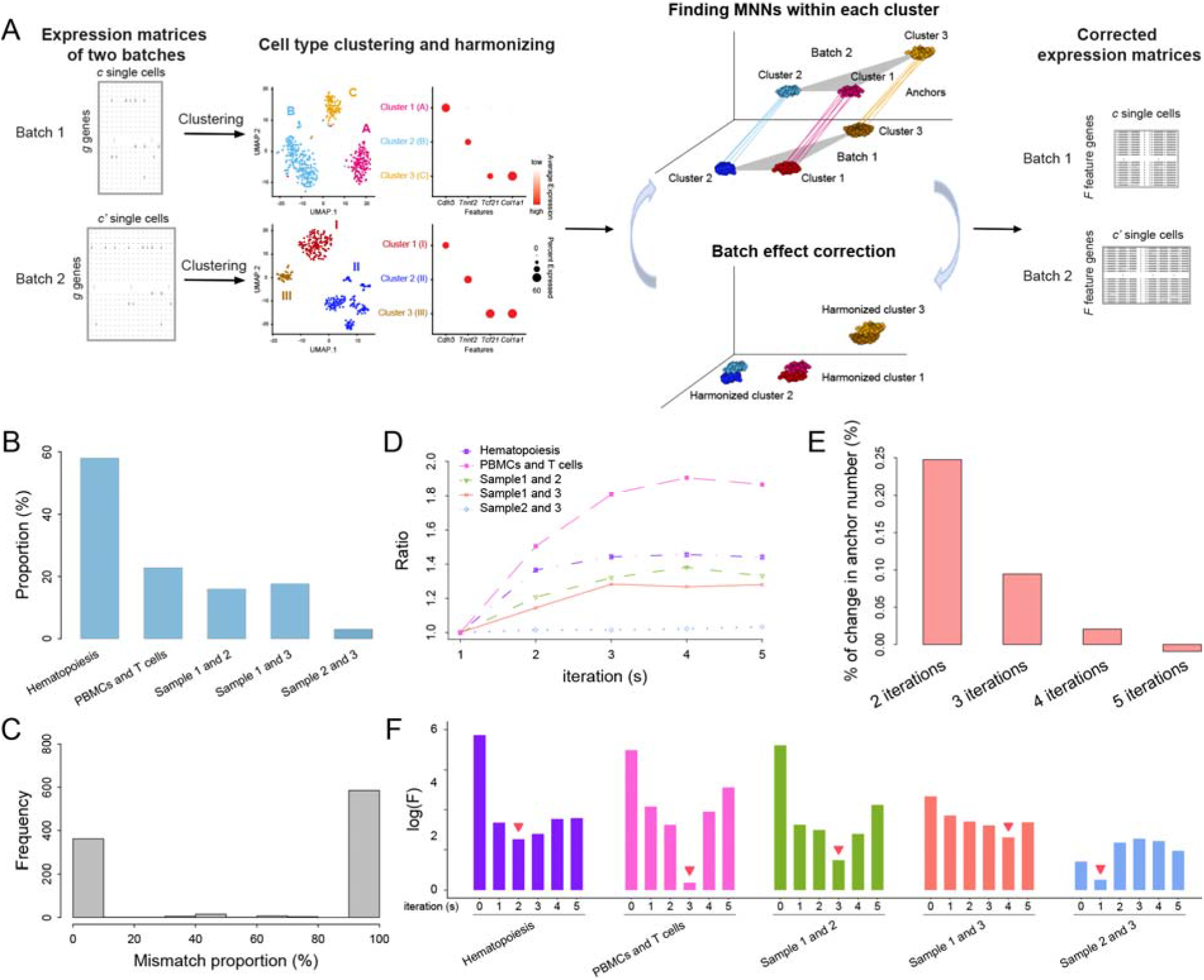
Overview of iSMNN. (**A**) Schematics of iSMNN. (**B**) Histogram of the proportion of mismatched mutual nearest neighbors (MNNs) (i.e., MNNs from a mismatched cell type) in the five sets of integration. (**C**) Histogram of the proportion of MNNs of a certain cell from a mismatching cell type in the hematopoietic datasets. (**D**) Ratio of the number of MNNs detected in each iteartion of batch effect correction, compared to the first iteration, in the five real datasets. (**E**) The average percentage changes in MNNs detected between the next two iterations of correction. (**F**) Logarithms of F-statistics for the corrected data after each iteration of correction in the five real datasets. Detailed information of the five real datasets is provided in Supplementary Table S1. Red arrow above indicates where best performance is attained.

To address this issue, we propose a new strategy that searches MNNs on the corrected data, where the systematic differences across batches are expected to be smaller than those in the original data, thus can improve the correction performance than only using the MNNs detected in the uncorrected data. According to our preliminary results, as the increase of iterations of MNN refining and batch effect correction, more MNNs were obtained across batches (**Figure 1D**). Specifically, an average of 24.8% more neighbors was archived when searching on the data after the first-iteration correction than on the original data (**Figure 1E**). In addition, with one more iteration of correction, the number of MNNs further increased by 9.5% on the top of that of the second iteration. However, there is no substantial changes in the MNN numbers in the forth and the fifth iteration between batches. More importantly, in two sets of real data, the optimal correction was achieved after a three-iteration correction, and in another two datasets, the best results were researched by two or four iterations of correction, respectively, depending on how large the batch effect is (**Figure 1F**).

According to these findings, here we present iSMNN, an iterative supervised batch effect correction method that performs multiple iterations of MNN refining and batch effect correction instead of one iteration correction with the MNN detected from the original expression matrix. With the further refined MNNs from corrected data, iSMNN would improve the correction accuracy compared to those using the one-iteration correction from the original data.

## RESULTS

### Overview of iSMNN

In the current implementation of iSMNN, we first input a harmonized label for each shared cell type across all the batches, either based on prior knowledge (e.g., known cell types and their corresponding marker genes) or inferred via unsupervised clustering followed by annotation of clusters within each batch, as described in SMNN [18]. With the harmonized cell type labels, iSMNN, following the Seurat v3 procedure (detailed in https://satijalab.org/seurat/v3.2/integration.html), selects top 2,000 most informative genes, following the default Seurat pipeline (detailed in **Supplementary Section 2**), across batches, and carries out dimensional reduction jointly across batches via diagonalized canonical correlation analysis (CCA). iSMNN then performs the first iteration of batch effect correction where MNNs are searched only within each matched cell type across batches. Batch effect correction is performed accordingly based on to the MNNs identified. To further improve the performance after the first iteration of correction, iSMNN implements multiple iterations of batch effect correction (**Figure 1A**). In each iteration, iSMNN matches MNNs of the same cell type across batches in the corrected results from the last iteration, and then refines the correction, with the updated MNN information. The performance after each iteration of correction, measured the degree of mixing for cells of the same cell type across batches, is quantified by F statistics using a two-way multivariate analysis of variance (MANOVA), where a smaller F value indicates a better mixing of cells across batches. The iterative MNN searching and batch effect correction continue until the F measure starts to increase. The correction results with the smallest F measure, deemed as the optimal results, are iSMNN output.

### Benchmarking in simulated data

We first assessed the performance of iSMNN using simulated data. One example is showed in **Figure 2** where the two batches before correction were completely separated from each other in the UMAP space (**Figure 2A; Supplementary Figure S2A**). After correction, all the three methods, iSMNN, MNNcorrect and Seurat v3 (denoted as “Seurat” for brevity unless version number specified otherwise) successfully mitigated the discrepancy between the two batches, especially for iSMNN that the cells of the same cell type were properly mixed across the two batches (**Figure 2D; Supplementary Figure S2D**). However, in Seurat corrected results (**Figure 2C; Supplementary Figure S2C**), there were still a large proportion of cells from cell types 2 and 3 of the second batch left unmixed with those from the first batch, and some cells from cell type 2 of the second batch (green unfilled triangles) were mixed with those from cell type 3 of the first batch (blue filled triangles). Moreover, across all the 30 simulations, iSMNN substantially reduced the F measure than Seurat and MNNcorrect (**Figure 2E**). Specifically, for the example we showed in **Figure 2**, the F value of iSMNN was 98.9% and 97.9% lower than that of Seurat and MNNcorrect, respectively (**Supplementary Figure S2E**). We further showed that iSMNN still outperforms Seurat and MNNcorrect under the scenarios where only partial cell type(s) are shared between batches or when the batches have a moderate or extreme difference in cell group composition (**Supplementary Figures S3 and 4**; detailed in **Supplementary Section 3**). These results suggest that iSMNN provides improved batch effect correction over alternative methods.

**Fig. 2.**
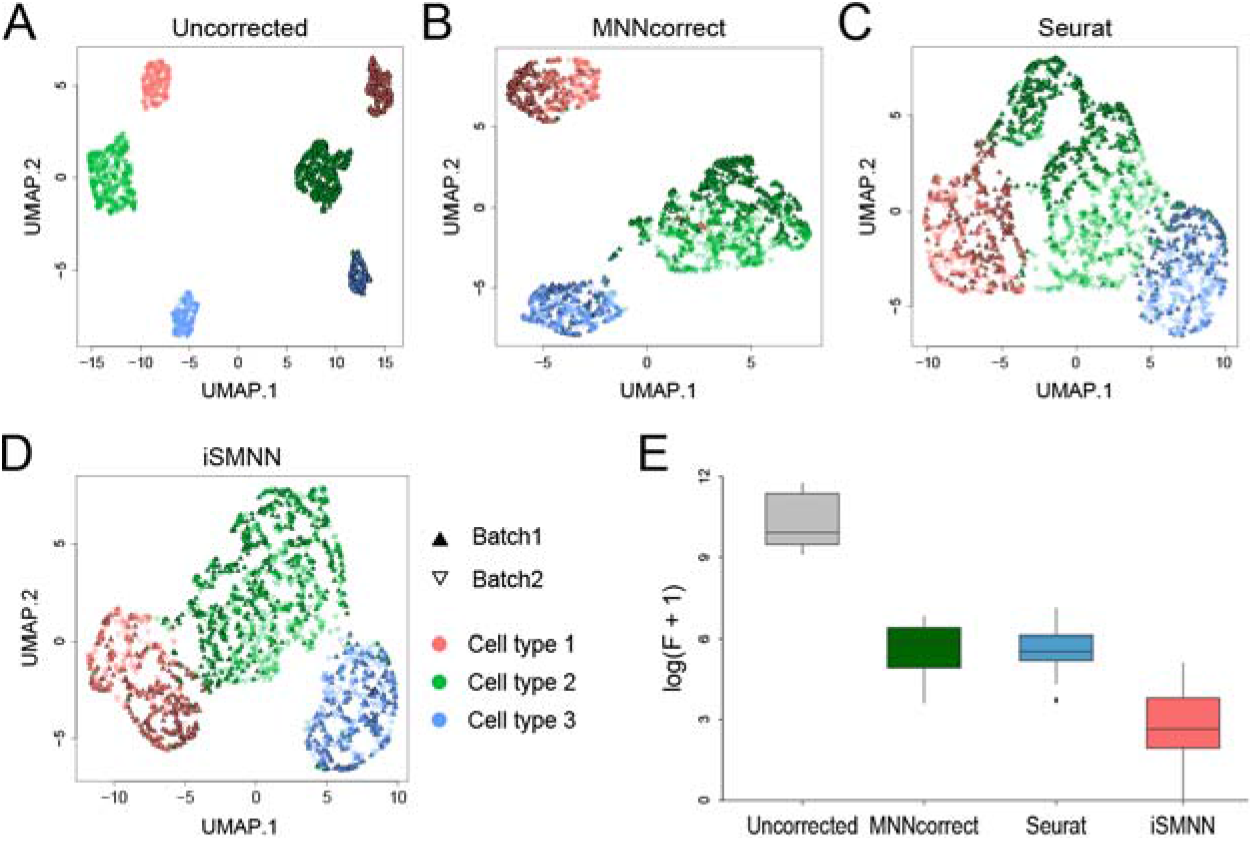
Performance comparison among iSMNN, Seurat and MNNcorrect in simulation data. (**A**), (**B**), (**C**) and (**D**) corresponds to the UMAP plots for the uncorrected, Seurat-, MNNcorrect and iSMNN-corrected results, respectively. (**E**) Boxplot of the logarithms of F-statistics for the merged data of the two batches before and after correction.

### Benchmarking in real data

We further evaluated iSMNN performance in real datasets (**Supplementary Table S1**), compared to six state-of-the-art methods (MNNcorrect, Seurat, Harmony [21], Scanorama [13], BBKNN [22] and SMNN [18]). Specifically, we used F statistic to rank the methods. The results showed that iSMNN consistently performed the best across all six sets of evaluations (**Figure 3**; **Supplementary Figures S5-9**), except for cardiac batch 2 and 3, where Harmony’s performance is slightly better than iSMNN and SMNN (**Supplementary Figure S8**). Regarding to the average ranking across different analysis, iSMNN outperforms all the alternative methods, followed by Harmony and Seurat.

**Fig. 3.**
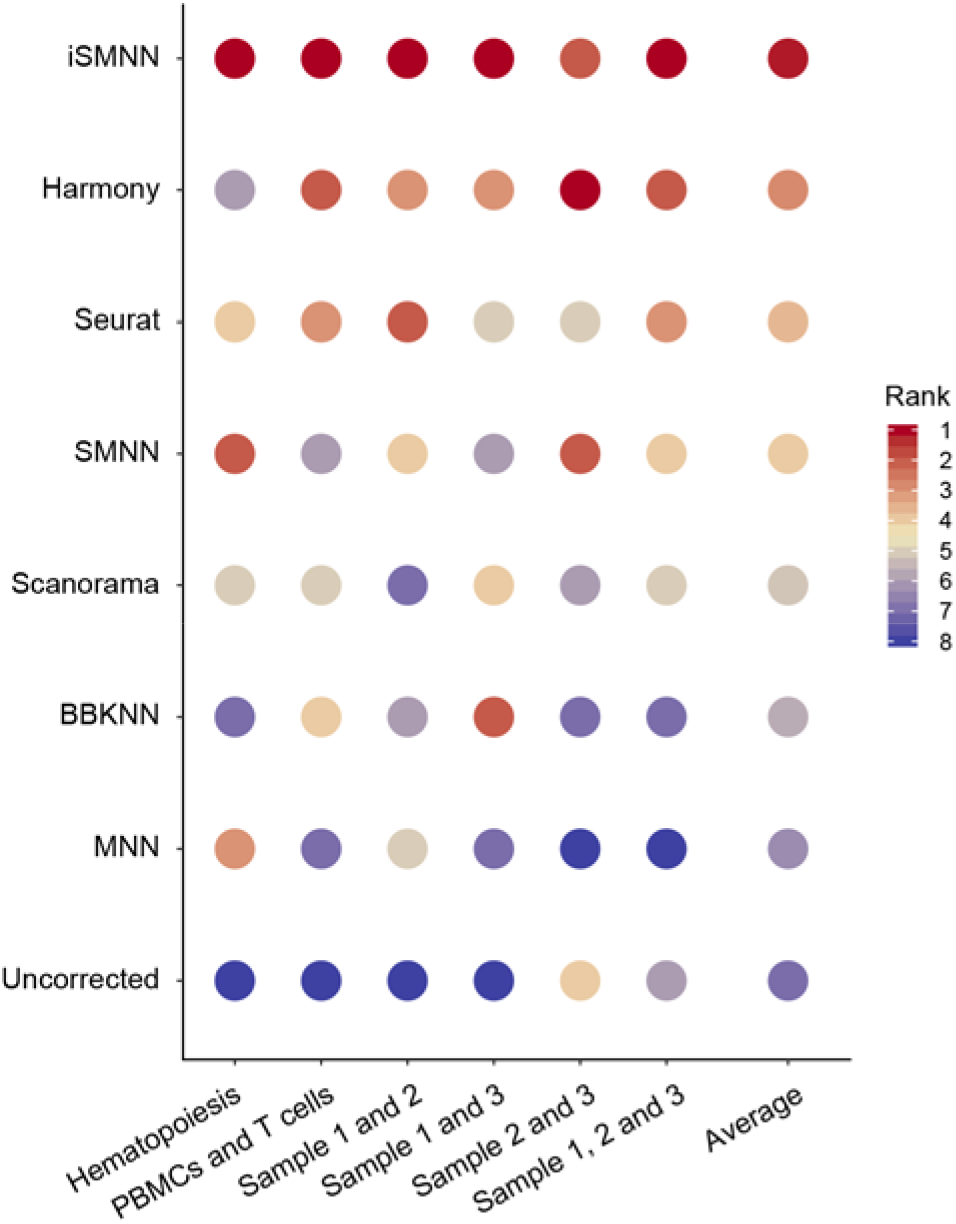
The rank of the seven batch effect correction methods based on their performance in benchmarking datasets measured by F-statistics. The methods are order according to the average rank across all datasets.

For the two hematopoietic datasets, compared to other methods, iSMNN can better merge cells across the two batches from the three shared cell types, namely common myeloid progenitor (CMP), granulocyte-monocyte progenitor (GMP) and megakaryocyte-erythrocyte progenitor (MEP) cells, as well as better distinguish the batch-specific cell types, such as multipotent progenitor (MPP) and multipotent long-term hematopoietic stem cells (LTHSC) cells, from the three shared cell types, than alternative methods (**Figure 4**). Consistently, F statistic of iSMNN was 56.9% - 99.1% lower than that of alternative methods (**Figure 4I**). For the datasets of human peripheral blood mononuclear cells (PBMCs) and T cells, iSMNN exhibited a better mixing of T cells from the two batches and smaller number of misclassified cells across different cell types (**Supplementary Figure S5**). These results suggest that iSMNN leads to improved batch effect correction.

**Fig. 4.**
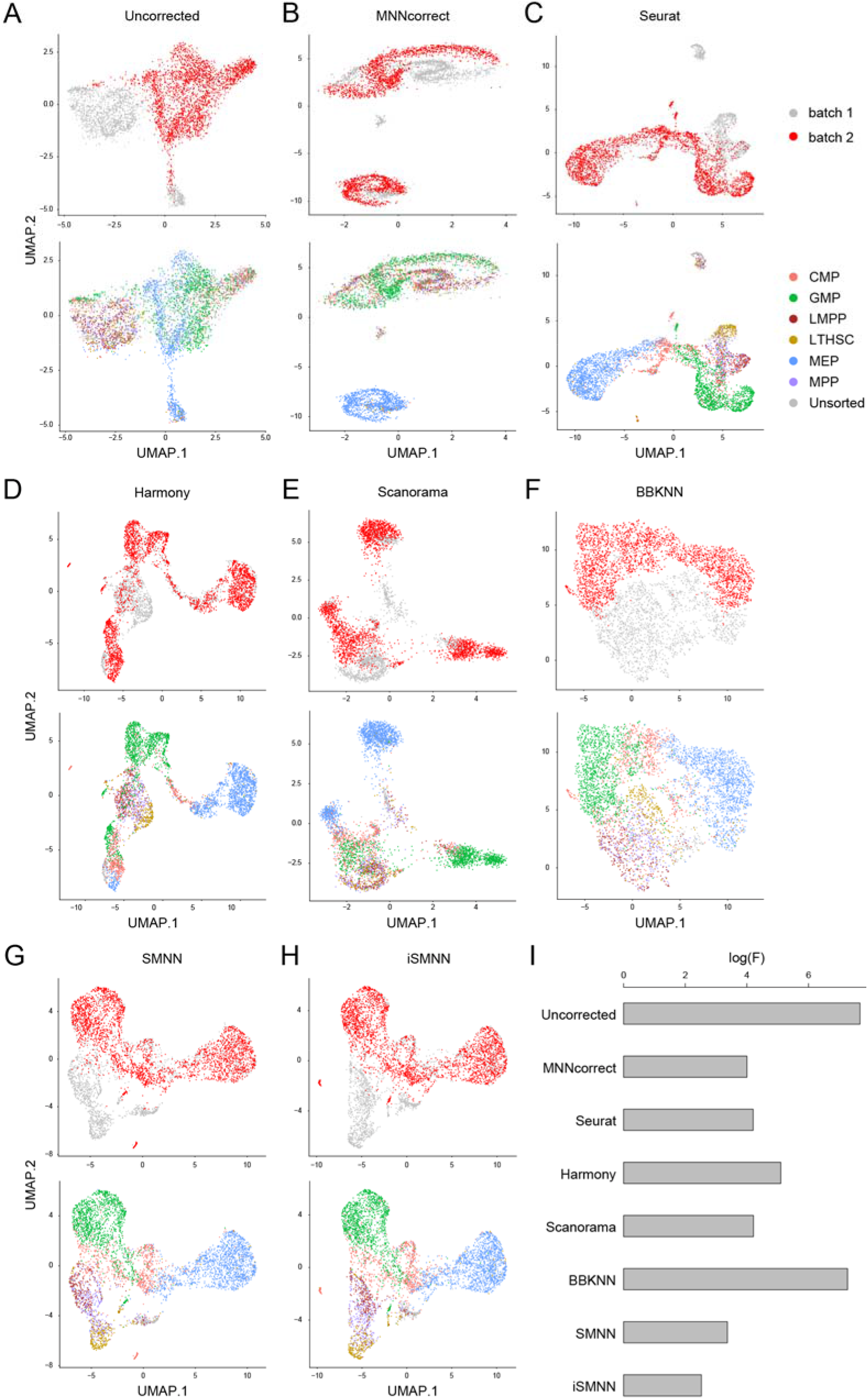
Performance comparison between iSMNN and alternative methods in two real datasets: the hematopoietic data. (**A-D**) UMAP plot for the (**A**) uncorrected, (**B**) MNNcorrect-, (**C**) Seurat-(**D**) Harmony-, (**E**) Scanorama-, (**F**) BBKNN-, (**G**) SMNN- and (**H**) iSMNN-corrected results in the hematopoietic data. (**I**) Logarithms of F-statistics for the merged data of the hematopoietic data before and after correction.

We further compared the performance between iSMNN and alternative correction methods in three cardiac cell datasets (see **MATERIALS AND METHODS** for details). In the batch we generated in this study (batch 1), we identified one group of CMs and five major cell types of non-myocytes (**Supplementary Figure S10**), where the five non-myocyte types were also present in the two batches from our previous study (batch 2 and 3), while the two CM groups were exclusively detected in batch 1. When we performed integrative analysis on batch 1 and 3 without batch effect correction, fibroblasts from the two batches were obviously separated in the UMAP space (**Figure 5A and E**). After batch effect correction, all the correction methods successfully brought cells from the two batches together (**Figure 5B-D**; **Supplementary Figure S7**). However, in the merged dataset, either Seurat and Harmony failed to distinguish cardiomyocytes (CMs) from endothelial cells (ECs) (**Figure 5F and G**). In contrast, results after iSMNN correction showed clear boundaries among different cell types (**Figure 4H**), and compared to alternative methods, iSMNN-corrected results exhibited smaller distance between two batches (**Supplementary Figure S7F**) and higher consistency within both CM and EC groups according to the average Silhouette Index (**Figure 5O**). We then compared the differentially expressed genes (DEGs) between CM and EC clusters identified by iSMNN and Seurat (**Figure 5I-N, Supplementary Figures S11-12).** Similar comparison with MNNcorrect can be found in **Supplementary Section 3 and Supplementary Figures S13.** Compared to ECs, a total of 143 and 126 DEGs were detected to be upregulated in CMs by iSMNN and Seurat, respectively, with 81 DEGs identified by both methods (**Figure 5I**). Expression profiling in batch 1, which was not affected by batch effect correction, showed that 51 out of 52 (98.1%) iSMNN-specific DEGs expressed in batch 1 indeed had higher expression levels in CMs than ECs, while only 77.6% (38 of 49) of the Seurat-specific DEGs was found to be upregulated in CMs (**Figure 5J and L**). In addition, Gene Ontology (GO) enrichment analysis revealed that the 62 iSMNN-specific DEGs are mainly involved in heart-related biological processes, such as ATP biosynthesis and metabolic processes, purine ribonucleotide metabolic process and calcium ion transmembrane transporter activity (**Figure 5M**). By contrast, Seurat-specific DEGs were found to be related to immune process, which does not seem relevant to the biological function of CMs (**Supplementary Figure S11**). To further validate DEGs specifically detected by iSMNN, we performed immunohistochemistry (IHC) for one CM-upregulated gene alpha B crystallin (*Cryab*) together with one CM-representative marker cardiac troponin T (*cTnT*) and EC-representative marker platelet endothelial cell adhesion molecule 1 (*Pecam1*). The staining results showed that anti-cTnT and anti-Pecam1 antibody can well demarcate CMs and ECs, respectively. Furthermore, Cryab+ cells are also positive for cTnT, but not positive for Pecam1 (**Figure 5P**), suggesting that *Cryab* is specifically expressed in CMs but not in ECs. Taken all together, our results indicate that iSMNN outperforms and retains more cell type specific features that are missed by alternative methods after correction.

**Fig. 5.**
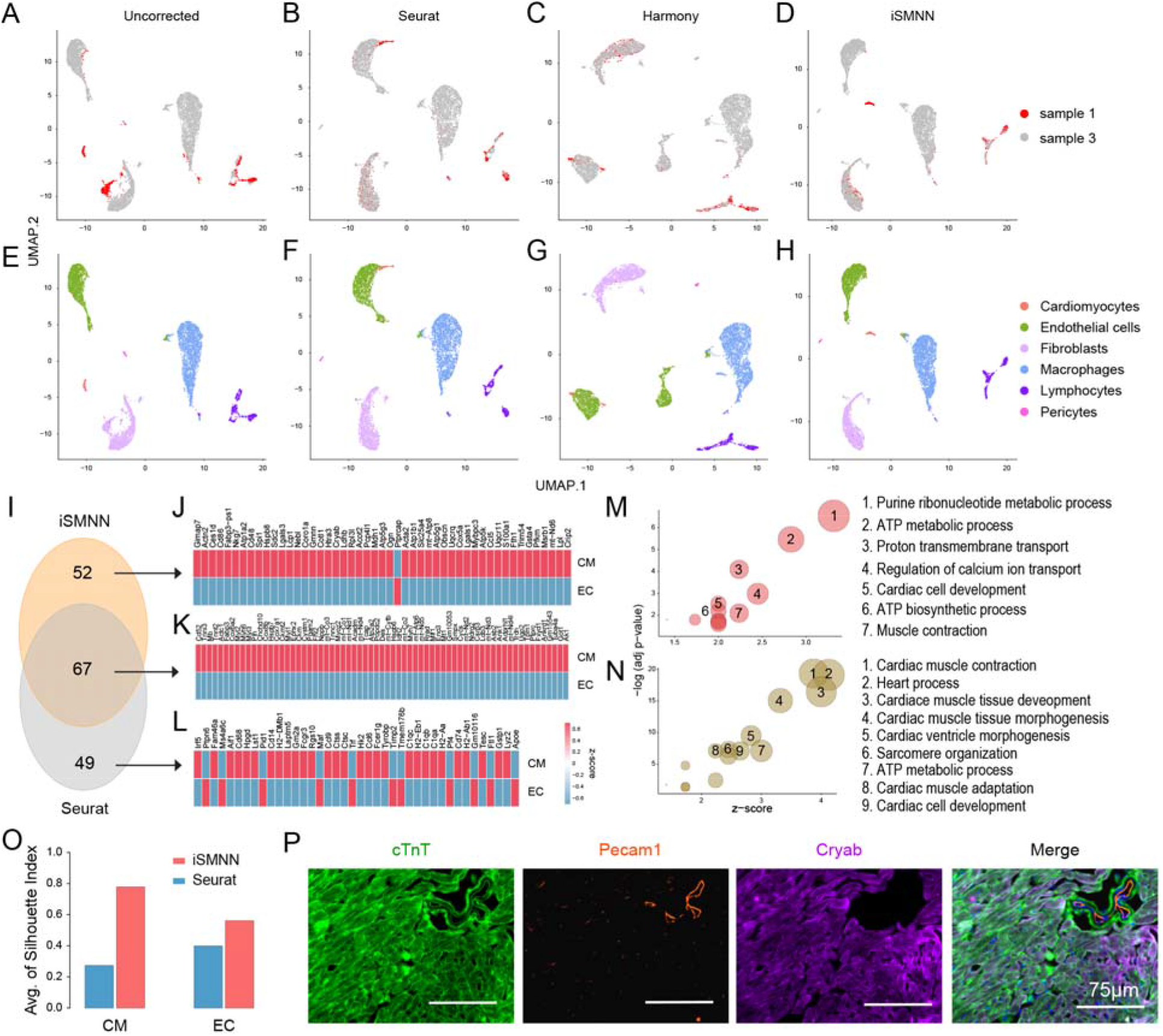
Performance comparison between iSMNN and alternative correction methods, MNNcorrect and Seurat, in two batches of cardiac data (batch 1 and 3). (**A**-**D**) UMAP plot for the (A) uncorrected, (**B**) Seurat-, (**C**) Harmony- and (**D**) iSMNN-corrected results for the two batches. (**E**-**H**) UMAP plots for the (**E**) uncorrected, (**F**) Seurat-, (**G**) Harmony- and (**H**) iSMNN-corrected results for the cell types across batches. (**I**) Overlap of differentially expressed genes (DEGs) upregulated in the cardiomyocyte (CM) cluster over the endothelial cell (EC) cluster after iSMNN and Seurat correction. (**J**-**L**) Heatmap showing gene expression profile of the DEGs upregulated in the CM cluster over the EC cluster, identified by (**J**) iSMNN specifically, (**K**) both iSMNN and Seurat and (**L**) Seurat specifically in cardiac batch 1. (**N**-**M**) Feature enriched GO terms for the overexpressed DEGs in CM cluster over EC cluster that were identified by (**N**) iSMNN specifically and (**M**) both iSMNN and Seurat. (**O**) Average Silhouette Index for the CM and EC clusters defined by iSMNN and Seurat, respectively. (**P**) Immunohistochemistry (IHC) staining for the typical CM marker cTnT, typical EC marker Pecam1, and one DEG Cryab specifically identified by iSMNN.

## DISCUSSION

In the current study, we present iSMNN, a supervised batch effect correction method for scRNA-seq data via multiple iterations of mutual nearest neighbor refinement. This work is built on our previously developed SMNN method, which has showed advantages in batch effect correction via supervised MNN detection over unsupervised correction methods. On top of SMNN, our iSMNN updates MNN detection iteratively and uses these MNNs for refined batch effect correction. Compared to the original data, systematic differences across batches after correction are much reduced, thus empowering the identification of more and better matched MNNs for improved batch effect correction in later iterations (**Figure 1D and E**). The procedure stops when the mixing performance of single cells of the same cell type across batches starts to deteriorate. This multiple-iteration approach substantially mitigates the MNN detection biases incurred by large batch effect between the original expression matrices and improves correction accuracy than a one-iteration approach (**Figure 1F**).

We benchmarked the performance of iSMNN with six alternative methods on three real scRNA-seq datasets. Our results clearly show that iSMNN can more effectively mitigate batch effect than alternative methods (**Figures 3-5**; **Supplementary Figures S5-9**). For example, our results for the two cardiac datasets (cardiac batch 1 and 3) showed that iSMNN substantially reduced differentiations across batches than MNNcorrect and Seurat (**Figure 5A-H**; **Supplementary Figure S7**), demonstrating that the advantages of iSMNN method in batch effect removal. More importantly, iSMNN better maintains cell-type distinguishing features. For the two cardiac datasets, iSMNN identified a more homogenous cluster for CMs than Seurat (**Figure 5H and O**), and iSMNN appears to more accurately recover features specific to each cell type, in terms of both gene expression profile and functional relevance. In particular, 98.1% of the CM-upregulated DEGs exclusively identified by iSMNN were observed to be more highly expressed in CMs than ECs, 26.4% improvement when compared to Seurat (**Figure 5J and L**). In addition, the CM-overexpressed DEGs specifically identified by iSMNN demonstrated biological function more relevant to CMs than those by Seurat (**Figure 5M**). IHC staining further validated that iSMNN-detected CM DEG *Cryab* was indeed specific to CM (**Figure 5P**). These results suggest that iSMNN can accurately maintain the cell-type specific features after batch effect correction than Seurat, which empowers valid downstream analysis and eliminates the spurious findings. Furthermore, although iSMNN performs supervised MNN detection, we show that it is be robust to the incompleteness of cell type annotation. Under the scenario where only partial cell type information is available, iSMNN still better mixed cells of the same cell type across batches than most of the alternative methods (**Supplementary Figure S14**; detailed in **Supplementary Section 3**).

As a multi-iteration procedure, iSMNN can potentially result in over-correction, with the performance deteriorating instead of improving in later iterations. One plausible reason is that as the iteration increases, the cells contributing to MNNs tend to concentrate disproportionately at certain areas (redder spots in **Supplementary Figure S15B and C**). Such disproportionate concentrations are not desirable because they may render the cells contributing to MNNs better mixed after correction, but the other cells not represented by MNNs will be “corrected” farther away from the MNNs selected, which will lead to an increased F-statistic and worse correction performance. Therefore, we select, as default, the first local minimum as the final corrected results of iSMNN. This default option is reasonable because in all our results, we find that once the F-statistic starts to increase, it never decreases below the first local minimum, suggesting the first local minimum is likely to well represent the global minimum. In addition, since the computational cost of iSMNN is not high (<20 minutes for a 10-iteration correction of two batches each containing 5000 cells; **Supplementary Figure S16**), we also provide users another two iteration options: in the first option, iSMNN runs for a fixed number of iterations (by default = 10), and takes the output with the lowest F statistic as the optimal correction results; in the second option, after the first local minimum is observed, an additional number of iterations (by default = 3) will be run to guard against further decrease of F-statistic after the first local minimal value.

In summary, leveraging iterative MNN refining, our iSMNN has demonstrated advantages in removing batch effect yet maximally retaining cell type specific biological features. We anticipate that our iSMNN will be a valuable method for integrating multiple scRNA-seq datasets, which can facilitate biological and medical studies at single-cell level.

## MATERIALS AND METHODS

### Simulation Framework

To assess the performance of iSMNN, we first performed simulation analysis following the framework described in Yang *et al*. [18]. Briefly, two batches X_*k*_ and Y_*l*_ were first simulated in a three-dimensional biological space following a Gaussian mixture model, where each component represents one cell type (Equations (1) and (2)).

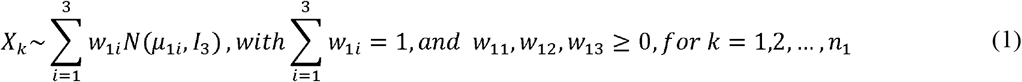

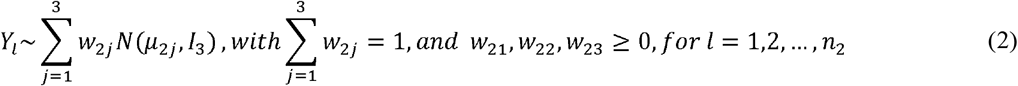

where *μ*_1*i*_is the three-dimensional vector specifying cell-type specific means for the *i*-th cell type in the first batch, reflecting the biological effect; similarly *μ*_2*i*_for the second batch; *n*_1_ and *n*_2_is the total number of cells in the first and second batch, respectively; *w*_1*i*_and *w*_2*i*_are the different mixing coefficients for the three cell types in the two batches; and *I*_3_ is the three dimensional identity matrix where diagonal entries are all ones and the rest entries are all zeros. In our simulations, we set *n*_1_ = 1000, *n*_2_ = 1100, (*w*_11_, *w*_12_, *w*_13_) = (0.3, 0.5, 0.2), and (*w*_21_, *w*_22_, *w*_23_) = (0.25, 0.5, 0.25.)

Then, batch effects were introduced to the three dimensional biological space, where the batch effect " was added to mean vectors of the three cell types in batch 1 to obtain the mean vectors of the three cell types for batch 2 (Equations (3)).

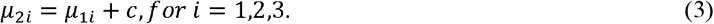

In this study, we set *c* = (0,5,2)^*T*^, (*μ*_11_, *μ*_12_, *μ*_13_) = ((5, 0, 0)^*T*^, (0, 0, 0)^*T*^, (0, 5, 0)^*T*^) and (*μ*_21_, *μ*_22_, *μ*_23_) = ((5, 5, 2)^*T*^, (0, 5, 2)^*T*^, (0, 10, 2)^*T*^). Finally, we projected the three-dimensional data with batch effect to the 2000dimensional gene expression space by linear transformation using the same random Gaussian matrix *P* within each batch (Equation (4) and (5)).

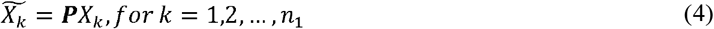

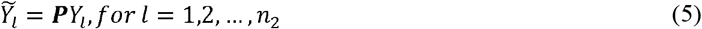

Here *P* is a 2000*×*3 Gaussian random matrix with each entry simulated from the standard normal distribution.

The simulation procedure was repeated 30 times with different random seeds. Both iSMNN and Seurat were applied to each set of simulated data. The merged results of the two batches before or after correction were visualized by UMAP [23], and the performance of each method was quantified by F statistics.

### Real data benchmarking

We also evaluated iSMNN’s performance in real datasets. iSMNN and six alternative correction methods: MNNcorrect, Seurat, Harmony, Scanorama, BBKNN and SMNN, were first applied to two published scRNA-seq datasets: 1) two hematopoietic samples generated using MARS-seq and SMART-seq2, respectively [24,25]; and 2) immune cells from human PBMCs and T cells from pancreas sequenced by 10X Chromium [2] (detailed in **Supplementary Table S1**). For both two datasets, cell type labels were assigned according to the annotations described in Haghverdi *et al.* [11], as well as the expression profile of the canonical markers. The performance of the three methods was compared by F statistics.

Furthermore, we applied iSMNN and alternative approaches on a scRNA-seq dataset for cardiac cells from adult murine heart (**Supplementary Table S1**). Specifically, we analyzed three batches of data: batch 1 that we have most recently generated scRNA-seq data for (NCBI Gene Expression Omnibus (GEO) accession number GSE161138; see **Supplementary Section 4** for experimental details), and batches 2 and 3, two other cardiac batches of adult murine hearts that we sequenced in our previous study (under review in *Cardiovascular Research*). Because the batch effect between batch 1 and 2/3 is more pronounced than that between batch 2 and 3, we assessed iSMNN, MNNcorrect and Seurat on three sets of integration under two scenarios: 1) batch 1 vs batch 2; 2) batch 1 vs batch 3; and 3) batch 2 vs batch 3. Given how the data for the three batches were generated, we anticipate 1) and 2) correspond to setting where batch effect is larger; and 3) where the batch effect is smaller. The performance was again measured by F statistics.

In additional, to measure how well the different cell types separate from each other after correction, we first performed unsupervised clustering on iSMNN, Seurat and MNNcorrect corrected datasets, respectively, using *FindClusters* function of Seurat [15] and assigned cell type label to each cluster according to the expression profiles of canonical markers (**Supplementary Table S2**). Then we carried out differential expression analysis between the clusters of CMs and ECs in iSMNN-, Seurat- and MNNcorrected-corrected datasets (detailed in **Supplementary Section 5**). Genes with a log(fold-change) > 0.25 and a adjusted *p*-value < 0.05 were considered as DEGs. GO enrichment analysis was carried out for three sets of DEGs: 1) those identified by both iSMNN and Seurat; 2) those identified exclusively by iSMNN; and 3) those detected exclusively by Seurat, respectively, using *clusterProfiler* [26]. To further validate whether the DEGs we identified in CMs are truly more highly expressed than in ECs, we implemented IHC staining for one DEG specifically identified by iSMNN, *Cryab*. Briefly, hearts of 3 month-old mice were sequentially perfused with 10mM KCl, PBS and perfusion buffer (0.5% PFA/5% Sucrose in PBS), then fixed in perfusion buffer overnight at 4℃. After dehydration in gradient concentration of sucrose, the heart was then embedded with OCT. Then the embedded blocks were sliced and cryosections were stored at −80℃. Before staining, sections (7 μm) were defrosted at room temperature for 5min. Then the sections were washed twice in PBST (PBS+0.1% Tween) and permeabilized with 0.2% Trition X-100 for 15min at RT. After permeabilization, sections were blocked with 5% BSA in PBS for 1 hour at RT, and then stained with primary antibodies against Cryab (Proteintech, 15808-1-AP, 1:200), cTnT (Sigma, MS-295-P, 1:200) and Pecam1 (BD, 550274, 1:50), in 1% BSA overnight at 4°C. The next day, after washing three times with PBS, sections were incubated with secondary antibody for 1 hour in the dark at RT followed by washing three additional times with PBS. Finally, the sections were mounted in Prolong Gold Antifade Mountant with DAPI (Invitrogen). The photos were taken using EVOS.

## DATA AND SOFTWARE AVAILABILIY

iSMNN is compiled as an R package, and freely available at https://github.com/yycunc/iSMNN and https://yunliweb.its.unc.edu/iSMNN. The data we adopted for benchmarking at from following: 1) two Mouse hematopoietic scRNA-seq datasets from Nestorowa *et al.* [24] (GEO accession number GSE81682) and Paul *et al.* [25] (GEO accession number GSE72857); 2) two 10X Genomics datasets of PBMCs and T cells from Zheng *et al.* [2] (https://support.10xgenomics.com/single-cell-gene-expression/datasets/); 3) three cardia datasets of adult murine hearts, one of which was generated in this study (GEO accession number GSE161138), and the other two are from Wang *et al.* (under review in *Cardiovascular Research*) (GEO accession number GSE157444).

## SUPPLEMENTARY DATA

Supplementary Data are available online.

## AUTHOR CONTRIBUTIONS

Y.L., L.Q., and Y.Y. initiated and designed the study. Y.Y., G.L. and Y.Y. implemented the model and performed simulation studies and benchmarking evaluation. Y.X., L.W. and J.L. performed experimental validation. Y.Y., G.L., L.Q., and Y.L. wrote the manuscript and all authors edited and revised the manuscript.

