## Supplementary file for "iSMNN: Batch Effect Correction for Single-cell RNA-seq data via Iterative Supervised Mutual Nearest Neighbor Refinement"

**Section 1. Correction for the cells in batch-specific clusters**

We first examined the cluster mismatching between batches. In the five sets of integrative analysis, 16.0 – 58.0% mutual nearest neighbors (MNNs) detected by Seurat v3 (denoted as “Seurat” for brevity unless version number specified otherwise) (1) were mismatched across cell types between the two batches (**Figure 1B**). Specifically, between the two hematopoietic datasets, only 36.9% (37.6%) of the cells have <10% (<50%) their nearest neighbors mismatched to different cell type(s) (**Figure 1C**). For the cardiac batch 2 and 3 that exhibit the smallest batch effect, there are still 3.6% (2.7%) of cells with >10% (>50%) of their nearest neighbors mismatched to different cell types (**Supplementary Figure S1**). These results suggest that mismatches are likely to be a big issue for the unsurprised MNN searching.

**Section 2. Informative gene selection for integration of batches**

In iSMNN, we select informative genes following the default Seurat pipeline, using Seurat’s *SelectIntegrationFeatures* function. Specifically, the top 2000 highly variable genes are first detected in each batch, and then the genes are ranked based on the number of times they are selected (as the top 2000) across batches (with the number ranging from 1 to the total number of batches). In order to break the ties (i.e., genes of the same rank or appearing the same number of times across batches), each gene is further ranked in each individual batch and re-ranked based on the median ranks across all batches. As the default, 2000 highest ranked genes across batches are selected as the informative genes for subsequent integration analysis.

**Section 3.** **Additional performance evaluation**

In this study, we further assess several other performance aspects of iSMNN. First, we evaluated the performance of iSMNN under the scenario that different batches have different cell population composition. To mimic the scenario where we only have partial cell type(s) shared between batches, we simulated two sets of data with only one or two of the three cell types shared. The results showed that, in both cases, iSMNN better mixed cells of the same cell type across batches than MNNcorrect and Seurat, reflected by the best/lowest F values from the iSMNN-correct results, compared with the corrected data by the other two methods (Supplementary Figure S3). Furthermore, we also simulated two sets of batches with a moderate or drastic difference in cell group composition. In the first dataset, there is a moderate difference where the average proportions of the three cell types in batch 1 are 29.8%, 50.0% and 20.2%; and 15.0%, 69.9% and 15.1% in batch 2. In the second dataset where the cell group compositions are drastically different, the average proportions of the three cell types are 29.6%, 50.3% and 20.1% in batch 1, and 5.1%, 90.0% and 4.9% in batch 2. In both two cases, iSMNN outperforms MNNcorrect and Seurat (Supplementary Figure S4). These results indicate that iSMNN is robust to differential cell population compositions across batches.

Secondly, we evaluated the robustness of iSMNN to the incompleteness of cell type annotation. Frist, we performed iSMNN correction by not feeding cluster labels for lymphocytes. The results showed good performance of iSMNN in mixing cells of the same cell type from different batches, although the performance is not as good as that with the labels from all four groups (**Supplementary Figure S14A-C**). Second, we randomly picked half of the cell cluster labels for iSMNN correction, to mimic the scenario where we only have partial prior knowledge regarding cell type annotations. The results showed that iSMNN with partial annotation information can still outperform all alternative methods except BBKNN (**Supplementary Figure S14D-F**). These results indicated that iSMNN is robust and performs reasonably well even with partial cell type annotation information.

To bias assess the robustness of our results to these criteria for differentially express gene (DEG) detection, we defined DEGs using different log-fold change cutoffs (0.1, 0.3 and 0.5). Our results showed that when the log-fold change cutoff increases, all the iSMNN-detected DEGs are exclusively upregulated in CMs, while 22.9% and 14.3% Seurat-detected DEGs are more highly expressed in ECs than CMs (**Supplementary Figure S12**), indicating a better correction performance of iSMNN than Seurat regardless of the log-fold change threshold. Moreover, we performed DEG comparison between iSMNN and MNNcorrect. Because of the larger number of DEGs identified by MNNcorrect than iSMNN, we ranked the CM-upregulated DEGs detected by MNNcorrect based on their adjusted p-value and took the top 119 DEGs (to match the number of DEGs detected in iSMNN corrected results) for downstream analysis. Of them, 32 DEGs were identified by both iSMNN and MNNcorrect (**Supplementary Figure S13**). The heatmap showed that, among the 88 MNNcorrect-specific CM-upregulated DEGs, the vast majority (84 or 95.5%) were indeed more highly expressed in CMs than ECs in batch 1. Results were even more consistent with iSMNN corrected results: 87 or 98.9% of the 88 DEGs showed consistent differential expression results in batch 1.

Finally, to assess the correction performance across iterations, we performed a real data-based simulation where we randomly split the cardiac sample 3 data from three major cell types (namely fibroblasts, endothelial cells and macrophages) into two pseudo-batches. Thus, the resulting two “pseudo-batches” should have no batch effects, by design. We then performed iSMNN correction on these two pseudo-batches using one to five iterations. The results (**Supplementary Figure S15**) showed that, F-statistic increases gradually from the first to the fifth iteration, and is larger/worse than that from the uncorrected data starting at the third iteration, suggesting that over-correction starts to manifest in later iterations. We examined the distribution of the cells identified as MNNs, and found that, as the iteration increases, the cells contributing to MNNs tend to concentrate disproportionately at certain areas (redder spots in **Supplementary Figure S15B and C**). Such undesirable disproportionate concentrations lead to an increased F-statistic and worse correction performance.

**Section 4. Single cell isolation and single-cell RNA sequencing**

Briefly, the whole heart was digested by gentle perfusing using Langendorff apparatus, and cardiac cells from ventricle were obtained. To balance the proportion of cardiomyocytes (CMs) and non-myocytes, the majority of CMs were depleted by low-speed centrifuge. Single live cells were enriched by flow cytometry and subjected to scRNA-seq using 10X Chromium platform. After quality control, a total of 947 cardiac cells were retained for downstream analysis. Unsupervised clustering was performed using Seurat v2 (2) following the default parameter settings, and cell type label was assigned to each cluster based on the expression pattern of canonical cell-type markers.

**Section 5. Detection of differentially expressed genes**

In this study, we define the DEGs from iSMNN and Seurat corrected results using *FindMarkers* function following Seurat’s default criteria. Specifically, a gene is considered a DEG if it has 1) a minimum fraction of 20% cells in either of the two cell types; 2) at least 0.25-fold higher expression (log-scale) in one group than the other; and 3) adjusted p-value from Wilcoxon Rank Sum test < 0.05. For the corrected results of MNNcorrect, the DEGs were defined with a minimum fraction cutoff of 20% and adjusted *p*-value cutoff of 0.05. Because of the larger number of DEGs identified by MNNcorrect than iSMNN, we ranked the CM-upregulated DEGs detected by MNNcorrect based on their adjusted *p*-value and took the top 119 DEGs (to match the number of DEGs detected in iSMNN corrected results) for downstream analysis.

**Supplementary Table S1.** Major characteristics of the three benchmarking datasets.

| **Dataset** | **Batch ID** | **Tissue origin** | **# of cells** | **Technical platform** | **Ref/GEO accession number** |
| --- | --- | --- | --- | --- | --- |
| Hematopoiesis | batch 1 | Mouse hematopoiesis | 1,920 | SMART-seq2 | (3) |
|  | batch 2 | Mouse hematopoiesis | 2,730 | MARS-seq | (4) |
| 10X Genomics | batch 1 | PBMC | 68,580 | 10X Genomics GemCode | (5) |
|  | batch 2 | T cells | 4,459 | 10X Genomics GemCode | (5) |
| Cardiac datasets | batch 1 | Mouse heart | 947 | 10X Genomics GemCode | GSE161138 |
|  | batch 2 | Mouse heart | 7,113 | 10X Genomics GemCode | GSE157444 |
|  | batch 3 | Mouse heart | 6,061 | 10X Genomics GemCode | GSE157444 |

**Supplementary Table S2.** Correspondence between known cell type annotations and clusters in iSMNN- and Seurat-corrected data.

**A.** iSMNN

| Cell type/Cluster | 0 | 1 | 2 | 3 | 4 | 5 | 6 | 7 | 8 | 9 | 10 | 11 | 12 | 13 |
| --- | --- | --- | --- | --- | --- | --- | --- | --- | --- | --- | --- | --- | --- | --- |
| Cardiomyocytes | 5 | 0 | 0 | 6 | 3 | 1 | 131 | 0 | 0 | 0 | 0 | 0 | 0 | 0 |
| Endothelial cells | 1 | 1571 | 0 | 4 | 0 | 0 | 1 | 120 | 63 | 0 | 2 | 1 | 53 | 2 |
| Fibroblasts | 1661 | 0 | 0 | 0 | 0 | 0 | 0 | 6 | 1 | 0 | 0 | 57 | 0 | 35 |
| Macrophages | 3 | 0 | 1553 | 948 | 0 | 1 | 0 | 0 | 52 | 0 | 20 | 1 | 0 | 0 |
| Lymphocytes | 1 | 0 | 4 | 6 | 329 | 167 | 0 | 0 | 0 | 1 | 43 | 0 | 0 | 0 |
| Pericytes | 1 | 1 | 0 | 0 | 0 | 0 | 0 | 0 | 0 | 66 | 0 | 0 | 0 | 0 |

**B.** Seurat

| Cell type/Cluster | 0 | 1 | 2 | 3 | 4 | 5 | 6 | 7 | 8 | 9 | 10 | 11 | 12 | 13 |
| --- | --- | --- | --- | --- | --- | --- | --- | --- | --- | --- | --- | --- | --- | --- |
| Cardiomyocytes | 6 | 2 | 1 | 4 | 3 | 0 | 129 | 0 | 1 | 0 | 0 | 0 | 0 | 0 |
| Endothelial cells | 1 | 1561 | 0 | 3 | 0 | 0 | 14 | 120 | 60 | 0 | 2 | 1 | 53 | 3 |
| Fibroblasts | 1663 | 1 | 0 | 0 | 0 | 0 | 1 | 6 | 0 | 0 | 0 | 57 | 0 | 32 |
| Macrophages | 3 | 0 | 1495 | 1004 | 0 | 1 | 1 | 0 | 52 | 0 | 21 | 1 | 0 | 0 |
| Lymphocytes | 1 | 0 | 3 | 8 | 332 | 164 | 0 | 0 | 0 | 0 | 43 | 0 | 0 | 0 |
| Pericytes | 1 | 0 | 0 | 0 | 0 | 0 | 0 | 0 | 0 | 67 | 0 | 0 | 0 | 0 |

**Supplementary Figures:**

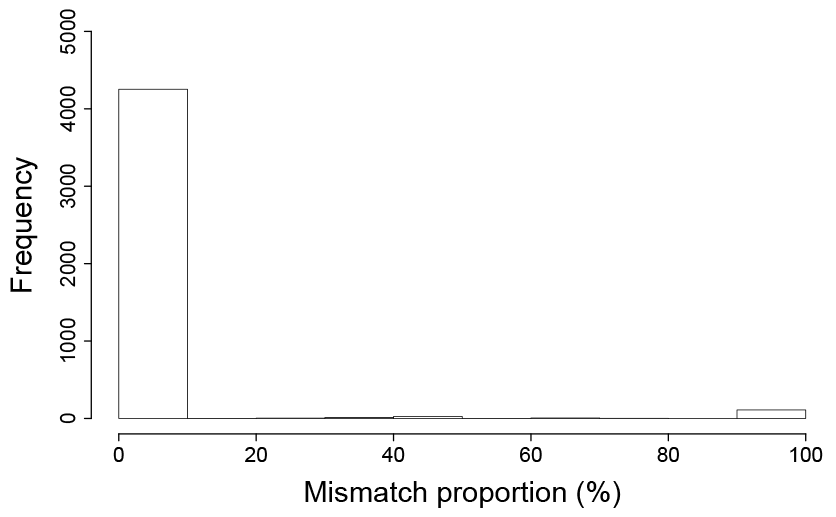

**Supplementary Figure S1.** Histogram of the proportion of MNNs of a certain cell from a mismatching cell type in cardiac batch 2 and 3.

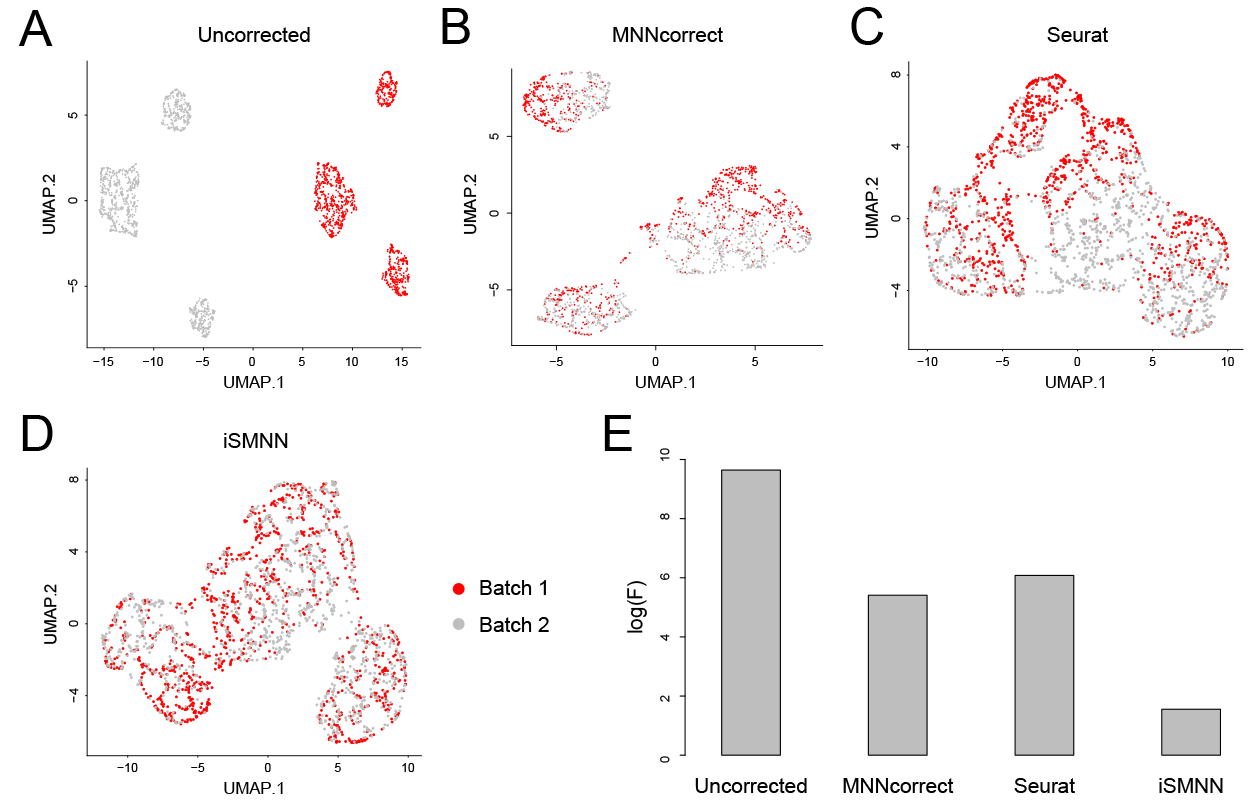

**Supplementary Figure S2.** Performance comparison among iSMNN, Seurat and MNNcorrect in simulated data. (**A**-**D**) UMAP plots for the (**A**) uncorrected, (**B**) Seurat-, (**C**) MNNcorrect and (**D**) iSMNN-corrected results for the two batches. (**E**) Logarithms of F-statistics for the merged data of the simulated data before and after correction.

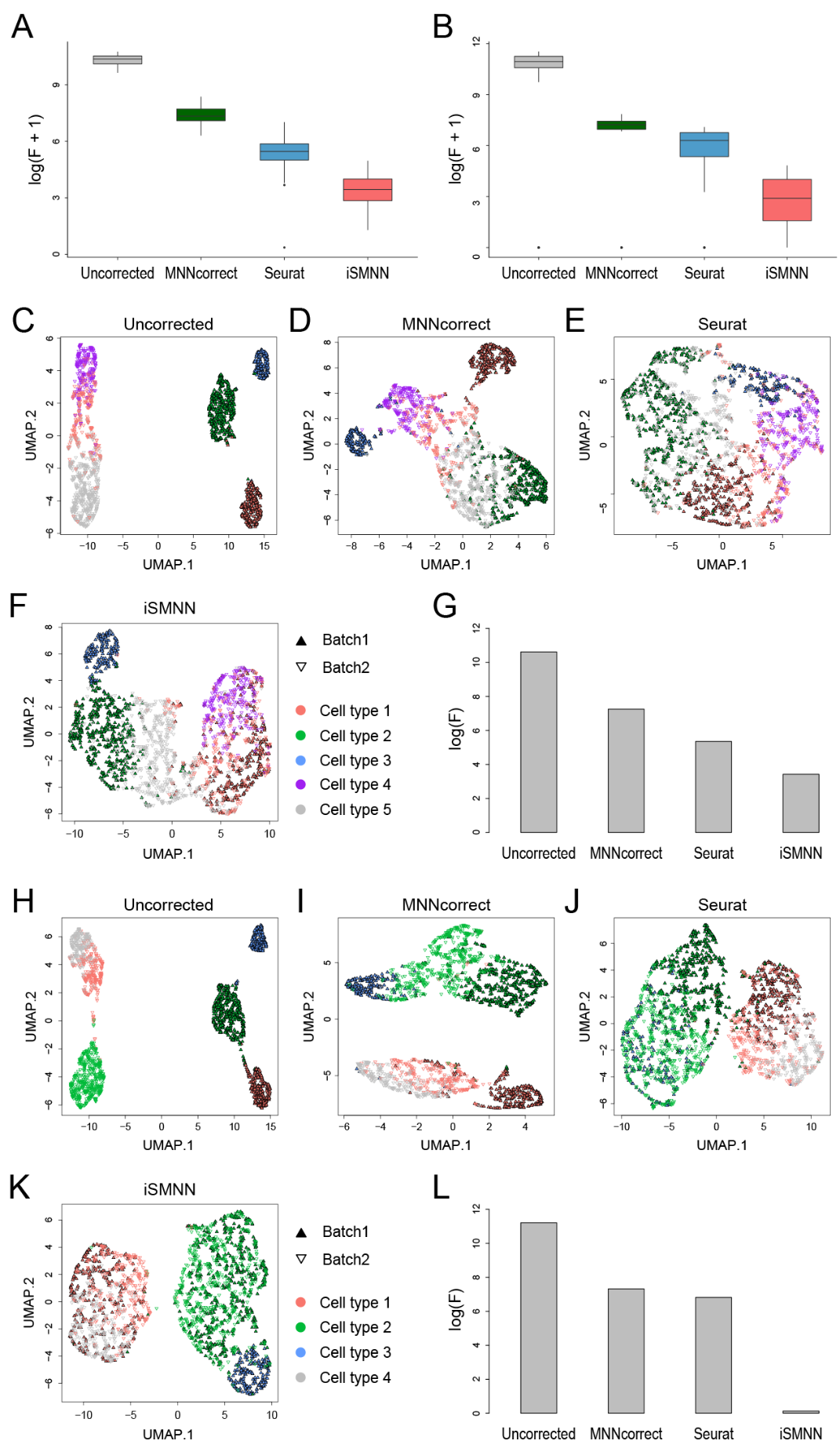

**Supplementary Figure S3.** Performance comparison between iSMNN and alternative correction methods, MNNcorrect and Seurat, based on all the cell types in two hematopoietic datasets. (**A**-**D**) UMAP plots for the (**A**) uncorrected, (**B**) MNNcorrect-, (**C**) Seurat- and (**D**) iSMNN-corrected results for all the cell types in two hematopoietic datasets. (**E**) Logarithms of F-statistics for the merged data of all the cell types in the two hematopoietic datasets before and after correction.

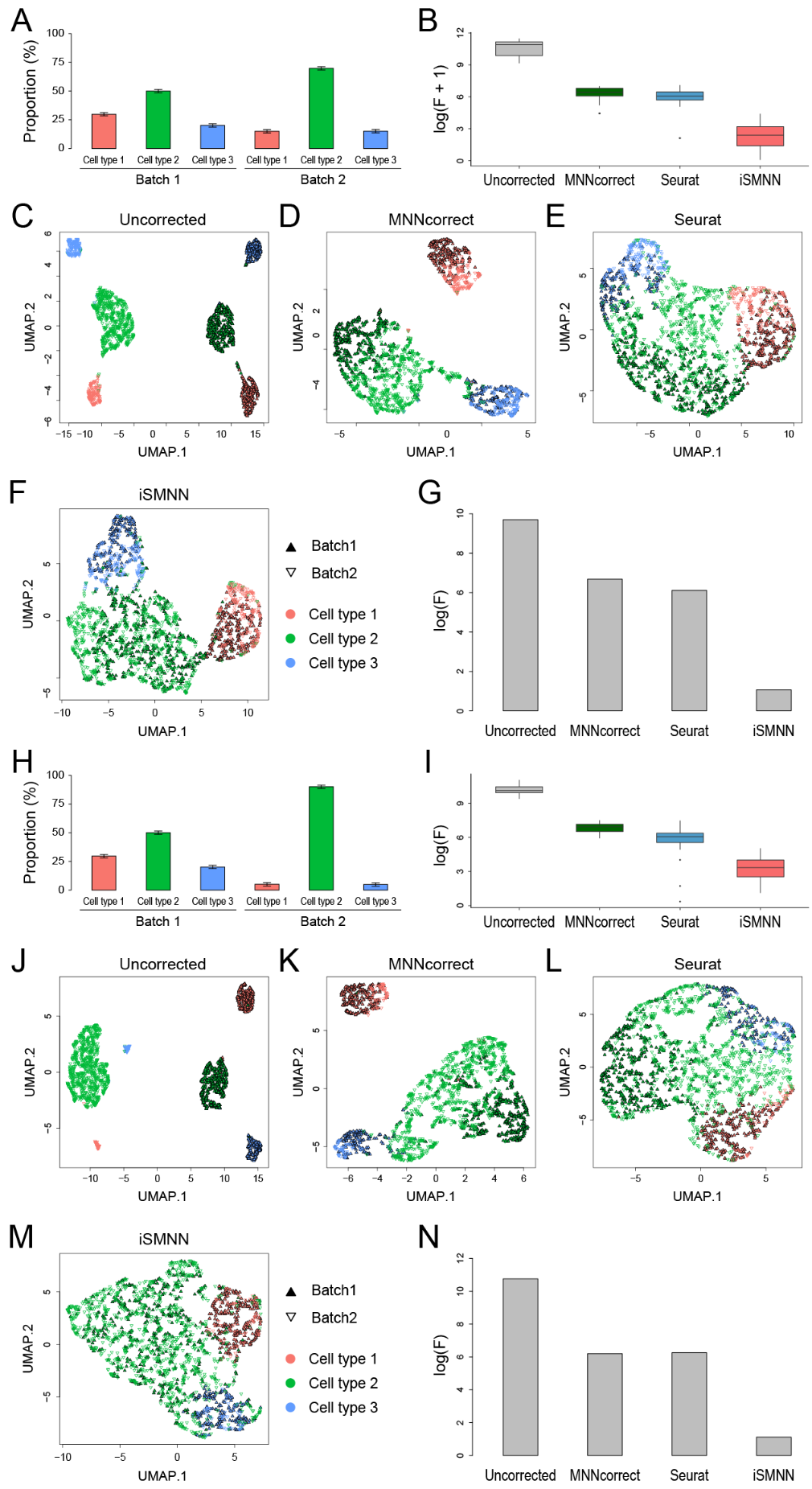

**Supplementary Figure S4.** Performance comparison between iSMNN and alternative correction methods, MNNcorrect and Seurat, based on all the cell types in two hematopoietic datasets. (**A**-**D**) UMAP plots for the (**A**) uncorrected, (**B**) MNNcorrect-, (**C**) Seurat- and (**D**) iSMNN-corrected results for all the cell types in two hematopoietic datasets. (**E**) Logarithms of F-statistics for the merged data of all the cell types in the two hematopoietic datasets before and after correction.

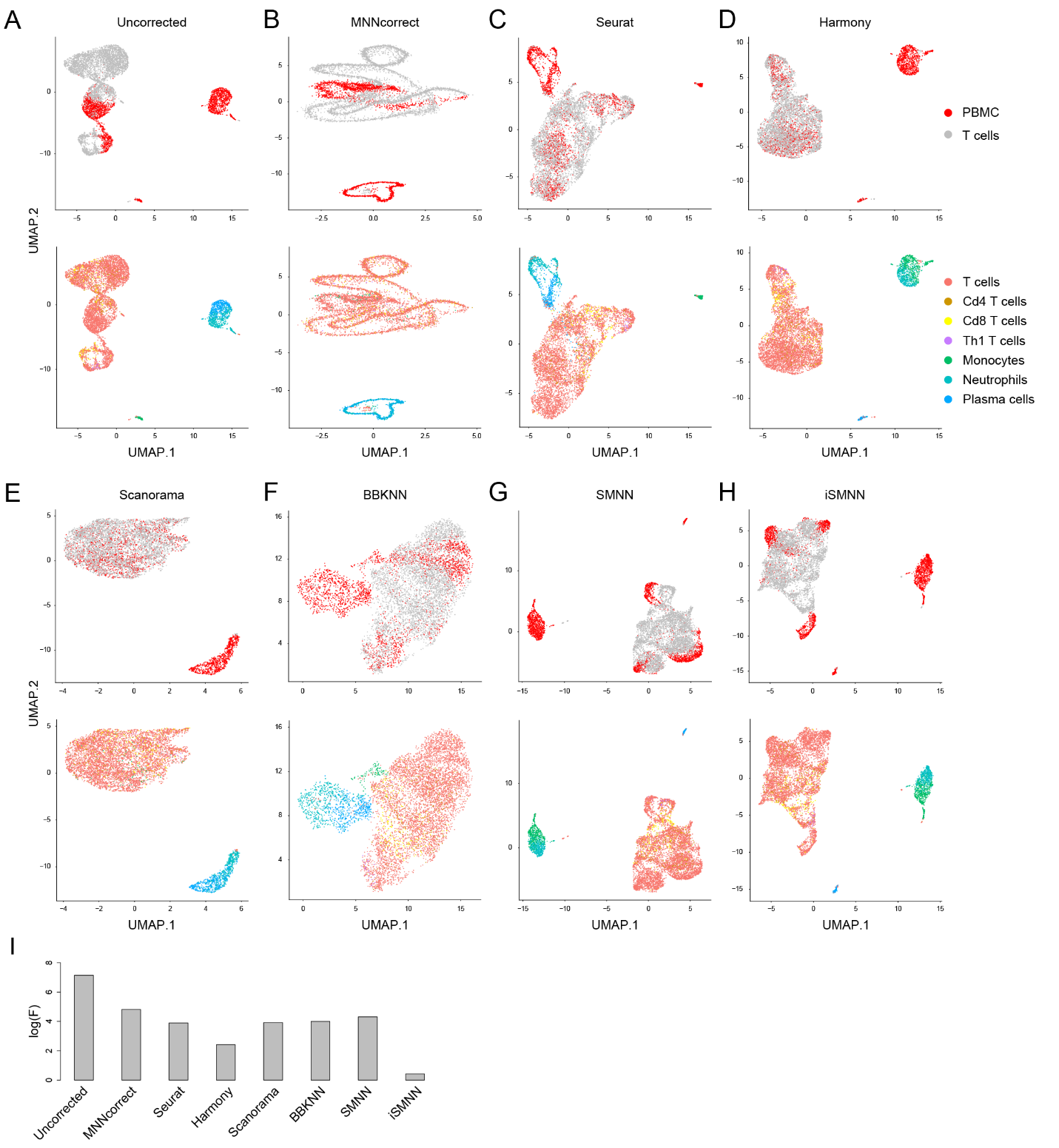

**Supplementary Figure S5.** Performance comparison between iSMNN and alternative correction methods in the datasets of human peripheral blood mononuclear cells (PBMCs) and T cells. (**A**-**D**) UMAP plots for the (**A**) uncorrected, (**B**) MNNcorrect-, (**C**) Seurat-, (**D**) Harmony-, (**E**) Scanorama-, (**F**) BBKNN-, (**G**) SMNN- and (**H**) iSMNN-corrected results for the two batches and the cell types across batches. (**I**) Logarithms of F-statistics for the merged data of all the cell types in the two batches before and after correction.

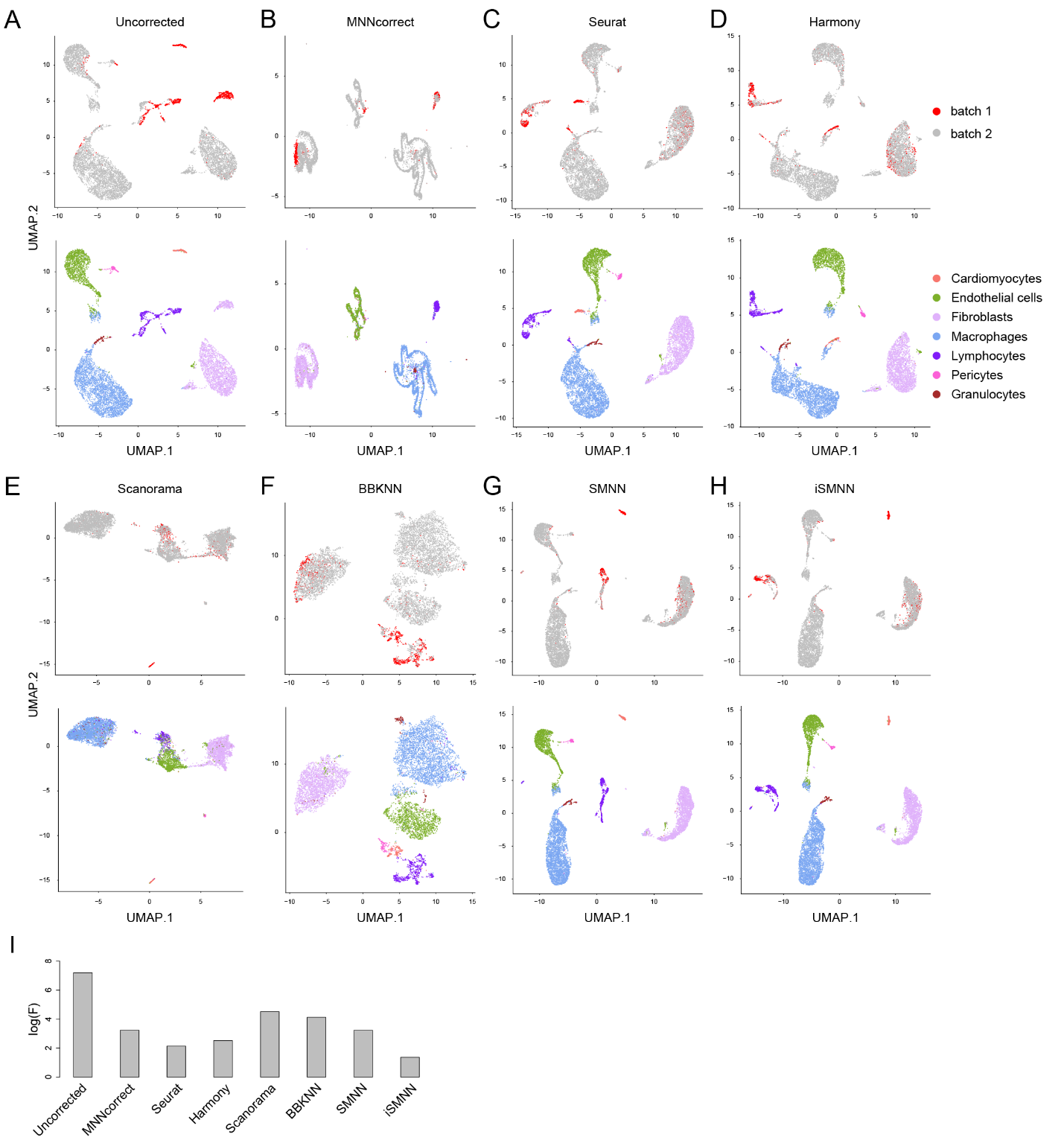

**Supplementary Figure S6.** Performance comparison between iSMNN and alternative correction methods in cardiac batch 1 and 2. (**A**-**D**) UMAP plots for the (**A**) uncorrected, (**B**) MNNcorrect-, (**C**) Seurat-, (**D**) Harmony-, (**E**) Scanorama-, (**F**) BBKNN-, (**G**) SMNN- and (**H**) iSMNN-corrected results for the two batches and the cell types across batches. (**I**) Logarithms of F-statistics for the merged data of all the cell types in the two batches before and after correction.

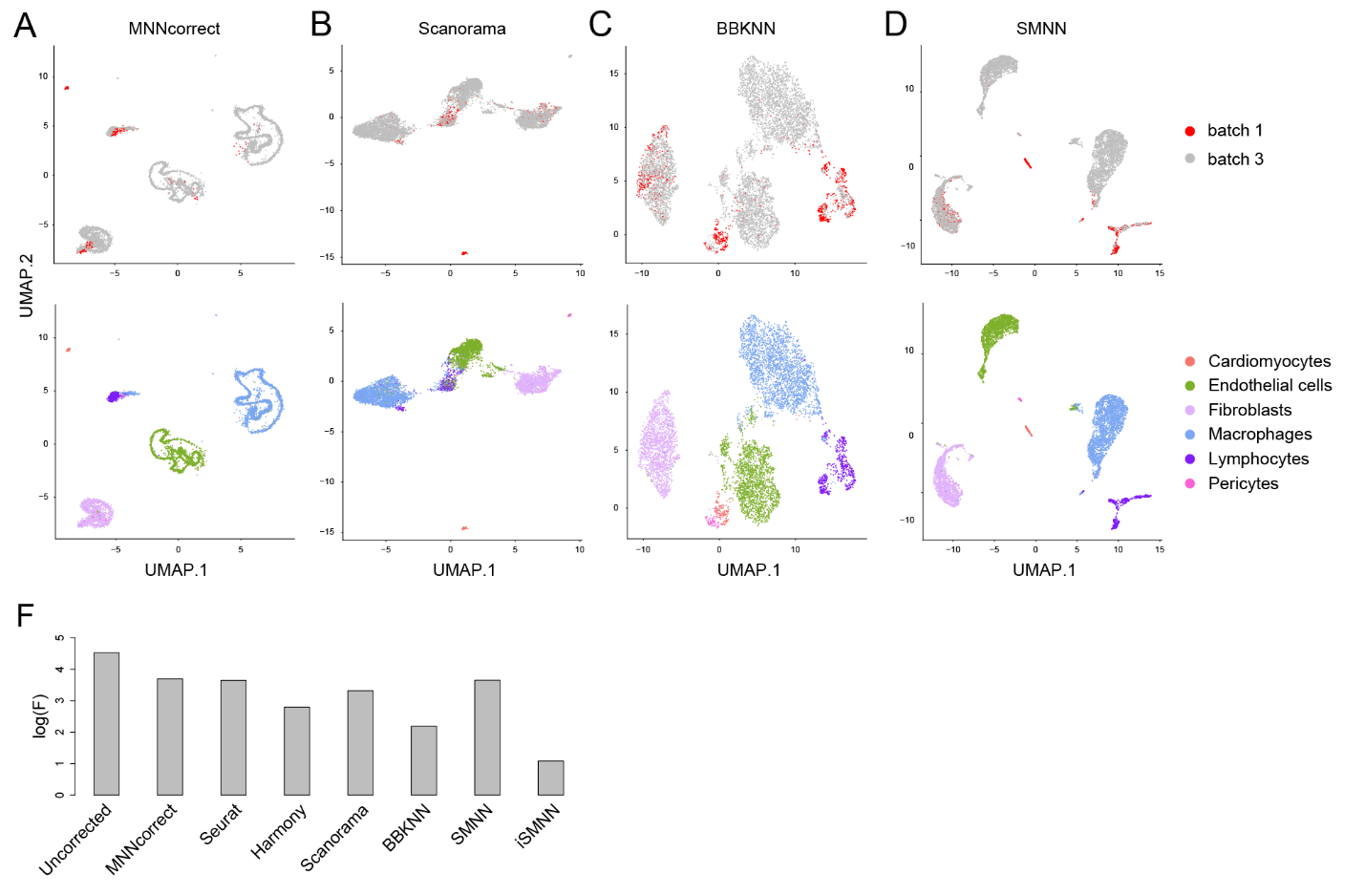

**Supplementary Figure S7.** Performance comparison between iSMNN and alternative correction methods, MNNcorrect, Scanorama, BBKNN and SMNN, in cardiac batch 1 and 3. (**A**-**D**) UMAP plots for the (**A**) uncorrected, (**B**) MNNcorrect-, (**C**) Seurat-, (**D**) Harmony-, (**E**) Scanorama-, (**F**) BBKNN-, (**G**) SMNN- and (**H**) iSMNN-corrected results for the two batches and the cell types across batches. (**I**) Logarithms of F-statistics for the merged data of all the cell types in the two batches before and after correction.

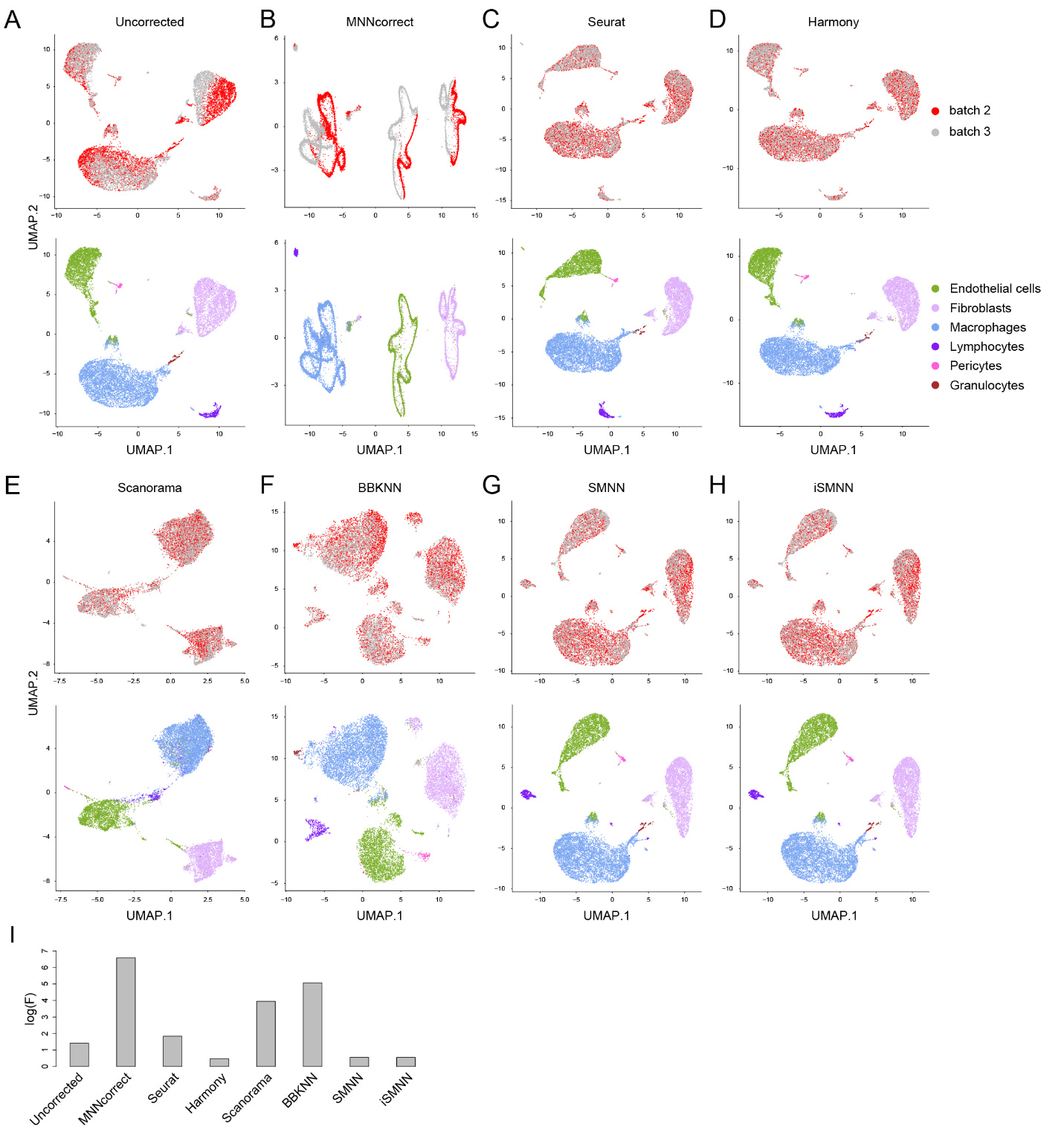

**Supplementary Figure S8.** Performance comparison between iSMNN and alternative correction methods in cardiac batch 2 and 3. (**A**-**D**) UMAP plots for the (**A**) uncorrected, (**B**) MNNcorrect-, (**C**) Seurat-, (**D**) Harmony-, (**E**) Scanorama-, (**F**) BBKNN-, (**G**) SMNN- and (**H**) iSMNN-corrected results for the two batches and the cell types across batches. (**I**) Logarithms of F-statistics for the merged data of all the cell types in the two batches before and after correction.

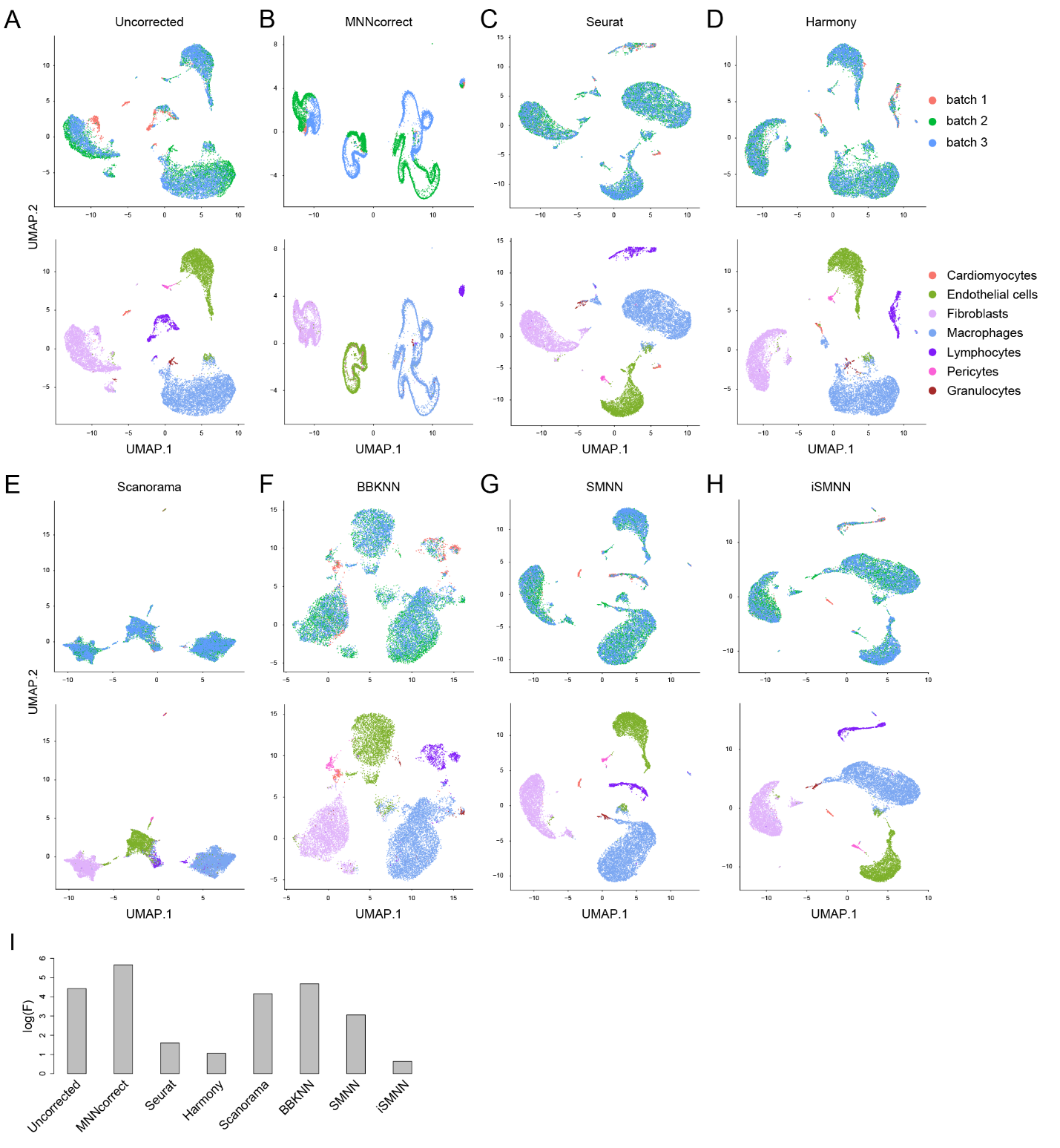

**Supplementary Figure S9.** Performance comparison between iSMNN and alternative correction methods in cardiac batch 1, 2 and 3. (**A**-**D**) UMAP plots for the (**A**) uncorrected, (**B**) MNNcorrect-, (**C**) Seurat-, (**D**) Harmony-, (**E**) Scanorama-, (**F**) BBKNN-, (**G**) SMNN- and (**H**) iSMNN-corrected results for the three batches and the cell types across batches. (**I**) Logarithms of F-statistics for the merged data of all the cell types in the three batches before and after correction.

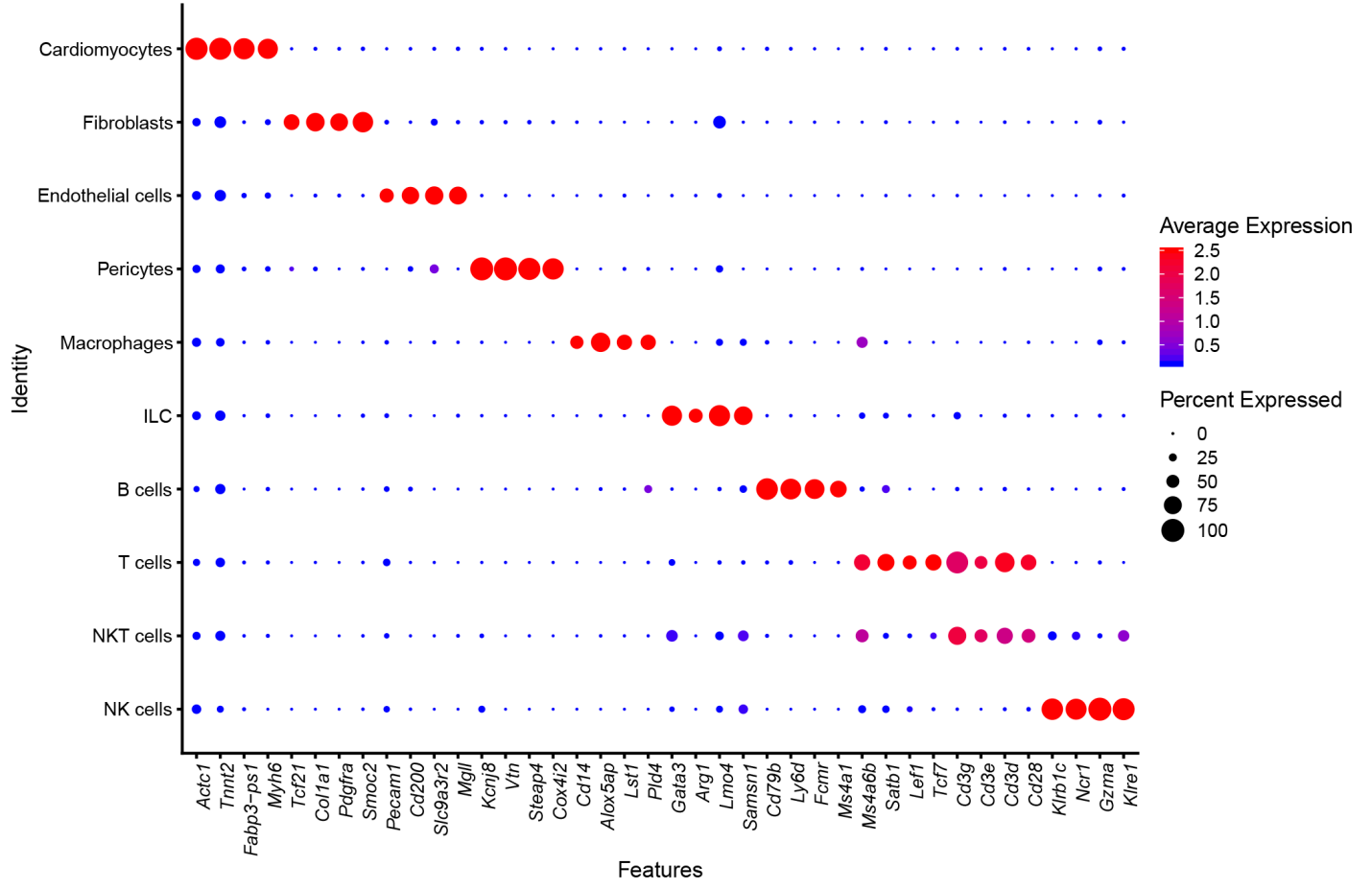

**Supplementary Figure S10.** Dotplot showing expression of the canonical markers for each cardiac cell type.

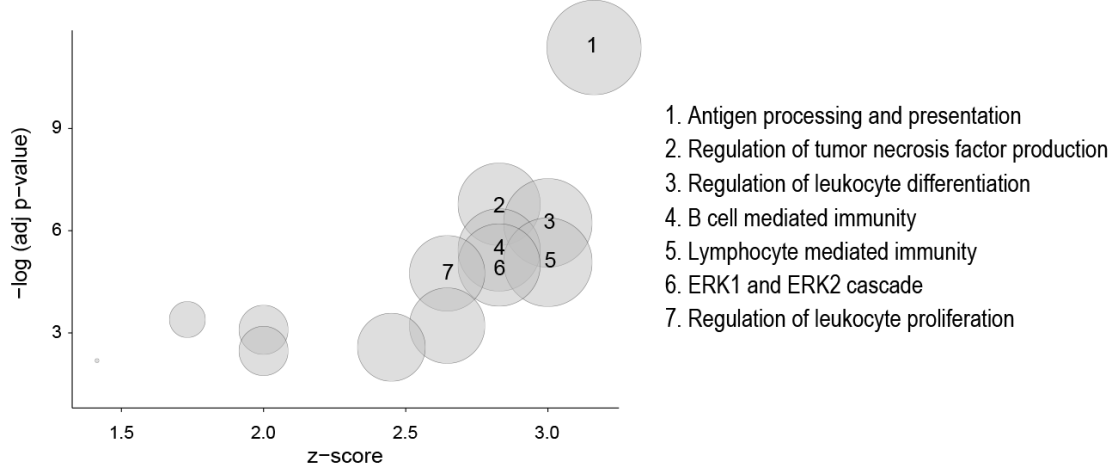

**Supplementary Figure S11.** Feature enriched Gene Ontology (GO) terms for the overexpressed differentially expressed genes (DEGs) in the CM cluster over the EC cluster that were specifically identified by Seurat.

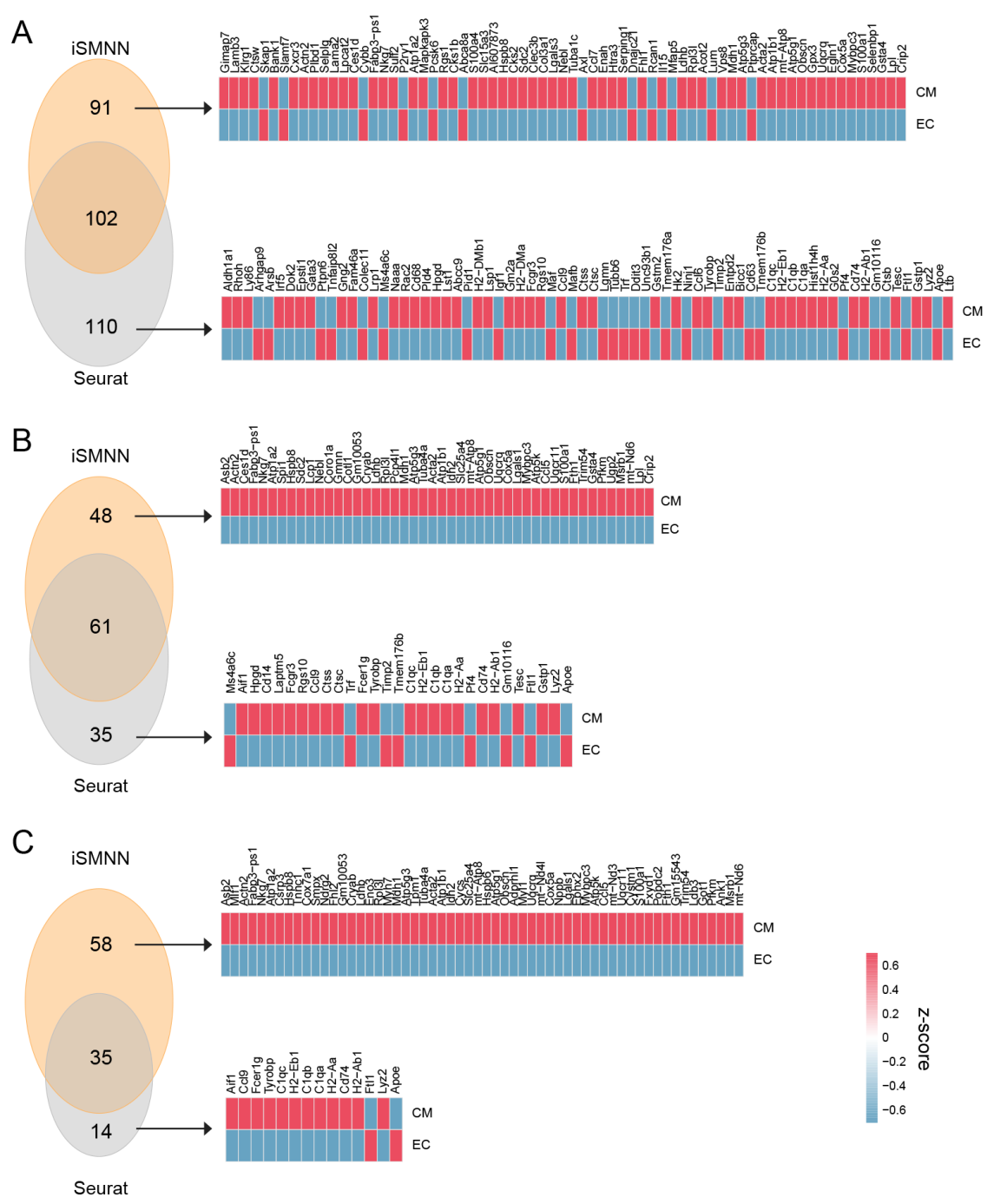

**Supplementary Figure S12**. Comparison of DEGs upregulated in the cardiomyocyte (CM) cluster over the endothelial cell (EC) cluster after iSMNN and Seurat correction using different log-fold change cutoffs, (**A**) 0.1, (**B**) 0.3 and (**C**) 0.5.

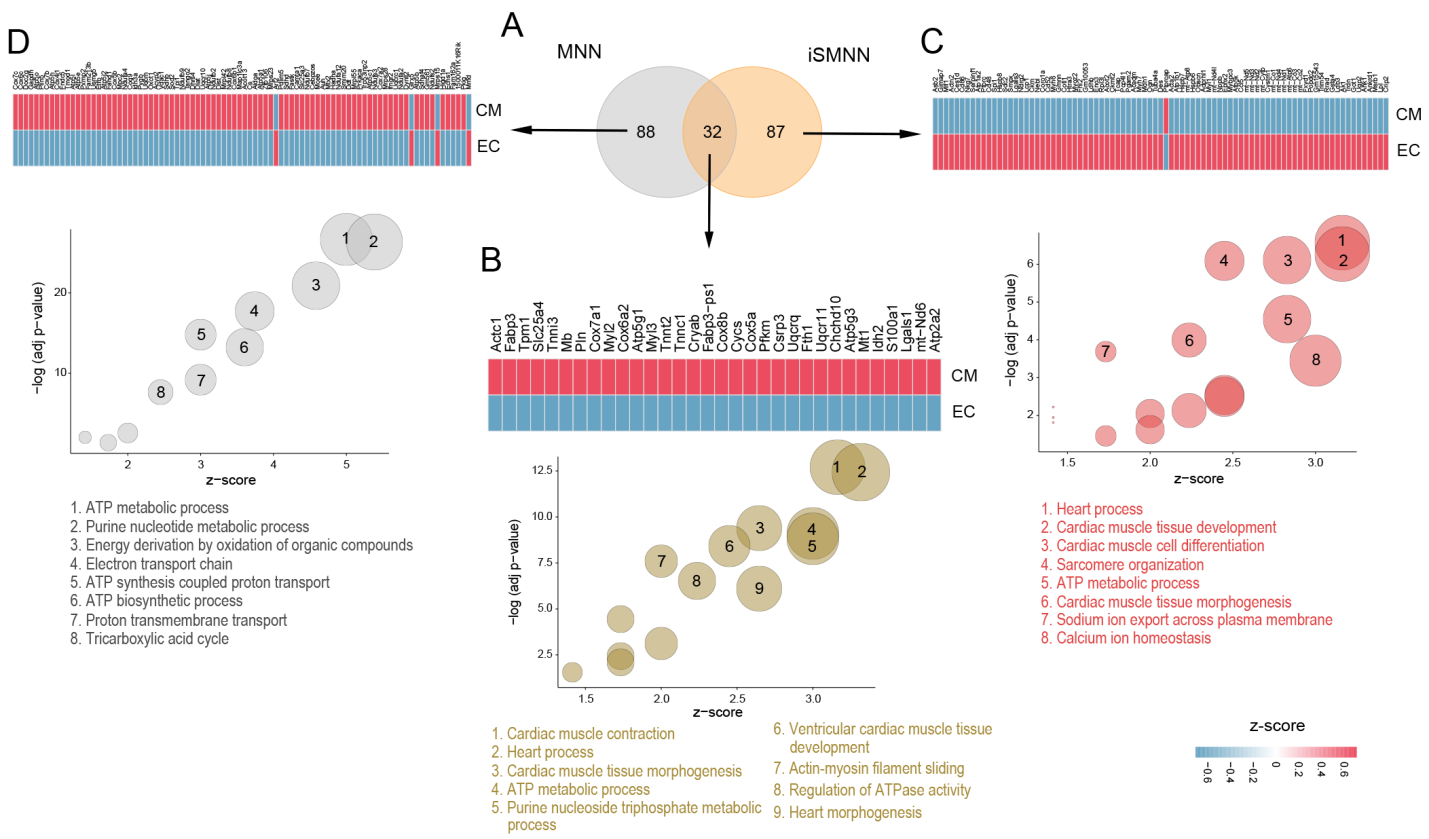

**Supplementary Figure S13.** Comparison of DEGs identified in the merged dataset by pooling the cardiac batch 1 and 3 after iSMNN and MNNcorrect correction. (**A**) Overlap of DEGs upregulated in CM cluster over EC cluster after iSMNN and MNNcorrect correction. (**B**-**D**) Feature enriched GO terms for the overexpressed DEGs in CM cluster over EC cluster that were identified (**B**) by both iSMNN and MNNcorrect, (**C**) specifically by iSMNN and (**D**) specifically by MNNcorrect in cardiac batch 1.

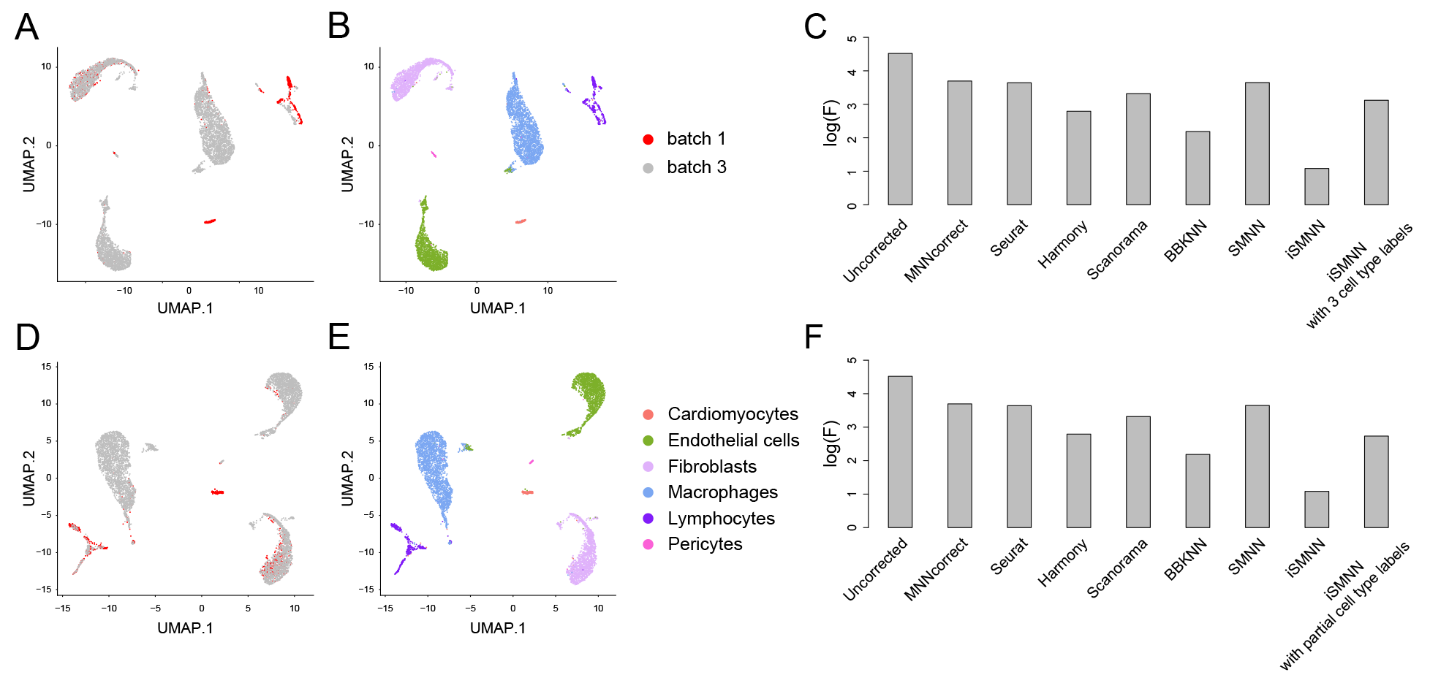

**Supplementary Figure S14**. Evaluation for the robustness of iSMNN to the incompleteness of cell type annotation. (**A**-**B**) UMAP plots for the iSMNN-corrected results by not feeding cluster labels for lymphocytes. (**C**) Logarithms of F-statistics for the merged data of all the cell types in the two batches before and after correction. (**D**-**E**) UMAP plots for the iSMNN-corrected results with randomly picked half of the cell cluster labels. (**F**) Logarithms of F-statistics for the merged data of all the cell types in the two batches before and after correction.

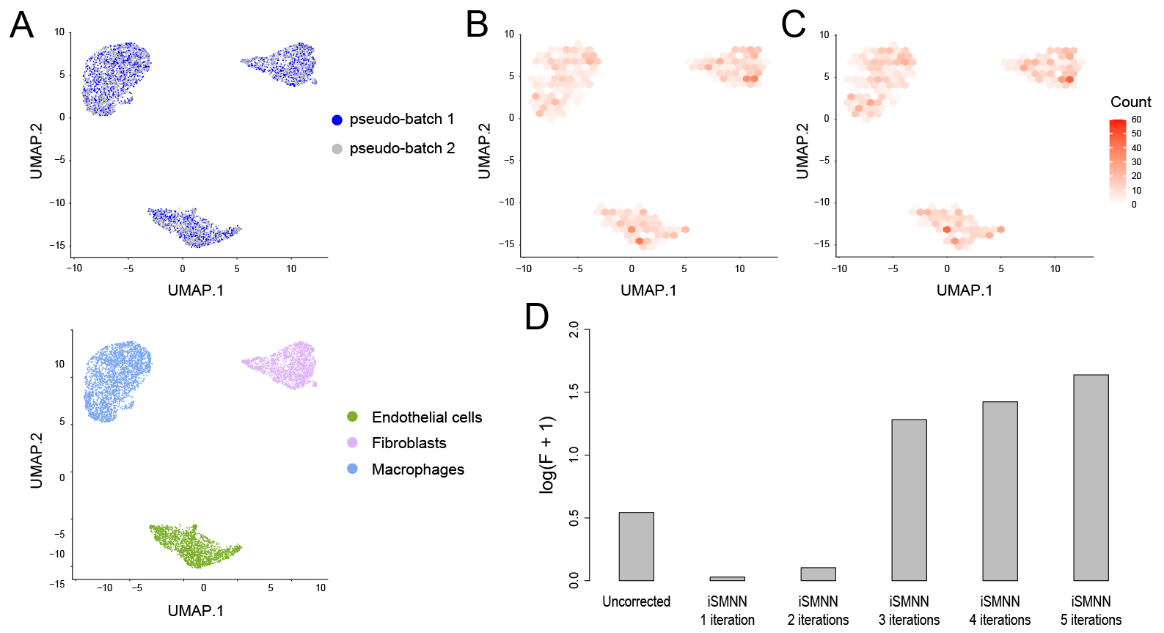

**Supplementary Figure S15**. Evaluation of the correction performance of iSMNN across iterations. (**A**) UMAP plots for the two pseudo-batches directly merged (without any batch effect correction). (**B-C**) The distribution of the cells identified as mutual nearest neighbors (MNNs) after correction with one iteration (**B**), or five iterations (**C**). (**D**) Logarithms of F-statistic for the merged data of the two batches before and after correction with one to five iterations.

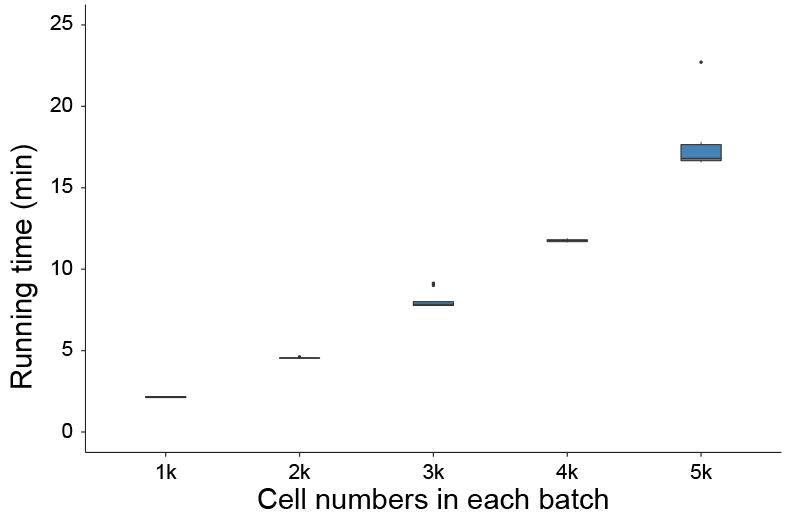

**Supplementary Figure S16**. Running time for two cardiac datasets, where each dataset contains 1,000 – 5,000 single cells. Running time (Y-axis) is in minutes (min). The computing time is from running on a PC with 2.90 GHz Intel Core i5-9400 CPU, 16 GB RAM and Windows 10 Pro operating system.
